# Evolution of Antibiotic Tolerance Shapes Resistance Development in Chronic *Pseudomonas aeruginosa* Infections

**DOI:** 10.1101/2020.10.23.352104

**Authors:** Isabella Santi, Pablo Manfredi, Enea Maffei, Adrian Egli, Urs Jenal

**Author notes:** These authors contributed equally to this work (Isabella Santi, Pablo Manfredi). **Author Contributions** I.S., P.M. and U.J. designed the study; I.S., P.M., E.M and A.E. collected and processed data; I.S. and P.M. performed the analyses; and I.S., P.M. and U.J. wrote the paper. **Competing interests** The authors declare no competing financial interests.

## Abstract

The widespread use of antibiotics promotes the evolution and dissemination of resistance and tolerance mechanisms. To assess the relevance of tolerance and its implications for resistance development, we used *in vitro* evolution and analyzed inpatient microevolution of *Pseudomonas aeruginosa*, an important human pathogen causing acute and chronic infections. We show that the development of tolerance precedes and promotes the acquisition of resistance *in vitro* and we present evidence that similar processes shape antibiotic exposure in human patients. Our data suggest that during chronic infections, *P. aeruginosa* first acquires moderate drug tolerance before following distinct evolutionary trajectories that lead to high-level multi-drug tolerance or to antibiotic resistance. Our studies propose that the development of antibiotic tolerance predisposes bacteria for the acquisition of resistance at early stages of infection and that both mechanisms independently promote bacterial survival during antibiotic treatment at later stages of chronic infections.

## Introduction

The great therapeutic achievements of antibiotics have been dramatically undercut by the steady evolution of survival strategies allowing bacteria to overcome antibiotic action ^1,2^. Although resistance plays a major role in antibiotic-treatment failure, bacteria can use resilience mechanisms such as tolerance to survive antibiotic treatment ^3^. Whereas experts and the public are well aware of problems related to increasing resistance, pathogen tolerance is not common knowledge, despite of being responsible for substantial morbidity and mortality ^4^. Resistance is generally drug specific and can be enhanced genetically through mutations modifying the drug target or by the acquisition of accessory genes such as efflux pumps ^5^. Such events lead to a decrease of the effective antimicrobial concentration and an increase of the minimum inhibitory concentration (MIC), which corresponds to the lowest drug concentration needed to prevent pathogen growth. In contrast, tolerance can lead to persistent infections despite a seemingly efficient treatment. In this case, a residual fraction of pathogens can resume growth after treatment was stopped, leading to infection relapses. For example, tolerance alters the kinetics of antibiotic killing without affecting MIC, leading to prolonged treatment necessary for pathogen eradication. Tolerance is thus measured by the minimum duration of killing of a specific fraction of the population ^6^. This phenotype has been related to non- or slow-growing bacteria that are able to survive bactericidal antibiotics for extended times ^7^. Tolerance can be adopted by all cells of a bacterial culture or only by a subpopulation called persisters ^8^. Bacterial populations with fractions of persisters are characterized by biphasic killing during treatment with bactericidal agents, where an initial rapid killing phase is followed by a phase of reduced killing ^7^.

Recent studies have shown that bacteria can rapidly evolve tolerance and persistence when exposed to antibiotics *in vitro* ^9–12^, suggesting that both represent successful strategies for bacteria to survive antibiotic treatment. In line with this, treatment efficacy during chronic infections was shown to be lost progressively without significant resistance development ^13,14^. Although challenging to diagnose ^15^, tolerant variants exist among environmental bacteria ^16^ and clinical isolates of human pathogens ^17–19^. Recently, it was proposed that antibiotic tolerance facilitates the evolution of drug resistance in laboratory conditions ^11,20^. However, it is still unclear if antibiotic tolerance plays a role in persistent infections and treatment failure ^14,19,21^ and if tolerance can facilitate resistance development in human patients ^10,15,20^.

Cystic fibrosis (CF) is the most common life-limiting, autosomal recessively inherited disease in Caucasian populations with the primary cause of death being respiratory failure resulting from chronic pulmonary infection ^22^. CF patients have reduced lung clearance capacity leading to the development of lifelong chronic infections caused by opportunistic bacterial pathogens such as *Pseudomonas aeruginosa*. Over time, treatment efficacy gradually declines and increasing inflammatory damage leads to fatal outcome ^23^. While infections are generally initiated by non-host adapted (i.e. naïve) strains found in the environment ^24–26^, *P. aeruginosa* undergoes significant microevolution during chronic infections of CF lungs ^13,27,28^. Despite recurrent application of high doses of antibiotics, a significant fraction of clinical isolates remains drug-sensitive ^13,14^. This argues that resistance development may not fully explain long-term survival of pathogens in the lung and that other strategies such as antibiotic tolerance contribute to the highly persistent nature of such infections.

Here we show that a substantial fraction of *P. aeruginosa* isolates from CF lungs has retained drug susceptibility, but has evolved various degrees of multi-drug tolerance. We demonstrate that recurrent exposure to high concentrations of antibiotics leads to the rapid development of tolerance, which generally precedes and boosts resistance development in *P. aeruginosa*. We show that tolerant strains display population heterogeneity with slow-growing subpopulations and that tolerance-mediated fitness costs can lead to the rapid loss of this phenotype after populations have acquired high-level resistance. Based on phenotypic and genotypic analysis of CF isolates, we propose that tolerance and resistance are alternative strategies contributing to *P. aeruginosa* persistence during long-term chronic infections.

## Results

### *P. aeruginosa* develops multi-drug tolerance during chronic infections of CF lungs

To explore the role of antibiotic tolerance in patients, we analyzed a large set of *P. aeruginosa* isolates (n=539) including strains from chronically infected CF patients (n=472), isolates from acute infections (n= 58 strains from 58 patients) and non-host adapted control strains from laboratory or environmental origins (n=9). CF patient isolates were sequentially isolated from 91 patients from five to 61 years old. For each strain we analyzed antibiotic resistance profiles (MIC) as well as the ability to survive exposure to tobramycin and ciprofloxacin over time (Fig. 1a; Extended data Fig. 1a,b). We chose these antibiotics because they have different modes of action and because they are used by clinicians to treat CF patients ^29^. Determination of MIC breakpoints according to clinical standards ^30^ revealed that clinically resistant strains were more common among isolates from chronic situations (22% tobramycin; 30% ciprofloxacin) compared to isolates from acute infections (6% tobramycin; 17% ciprofloxacin). When analyzing the survival of isolates with MIC values below the clinical breakpoint ^30^ during exposure to high concentrations of tobramycin or ciprofloxacin we observed large differences in killing kinetics within groups of isolates with identical levels of resistance (Fig. 1a; Extended data Fig. 1a,b). In particular, most isolates from chronic infections showed strongly enhanced tolerance and persistence, whereas isolates from acute infections were rapidly killed upon drug exposure (Fig. 1a,b; Extended data Fig. 1a-d). Minimum duration for killing of 99% (MDK99) and 99.99% (MDK99.99) of the bacterial population were proposed as measures for tolerance and persistence, respectively ^6^. Because of the high survival profile of several clinical isolates MDK99.99 was not a useful measure for persistence, we decided to quantify persistence as the level of survival after 7 hrs treatment. To clearly the two phenomena, tolerance was quantified as the proportion of surviving bacteria after 1 h of treatment, a value that correlates well with other measures for tolerance including the minimum duration for killing of 99% of the population (MDK99) ^6^ (Extended data Fig. 1e). Importantly, most antibiotic-sensitive isolates from CF airways showed increased survival in the presence of both drugs (Fig. 1d), emphasizing the multi-drug nature of tolerance.

**Figure 1:**
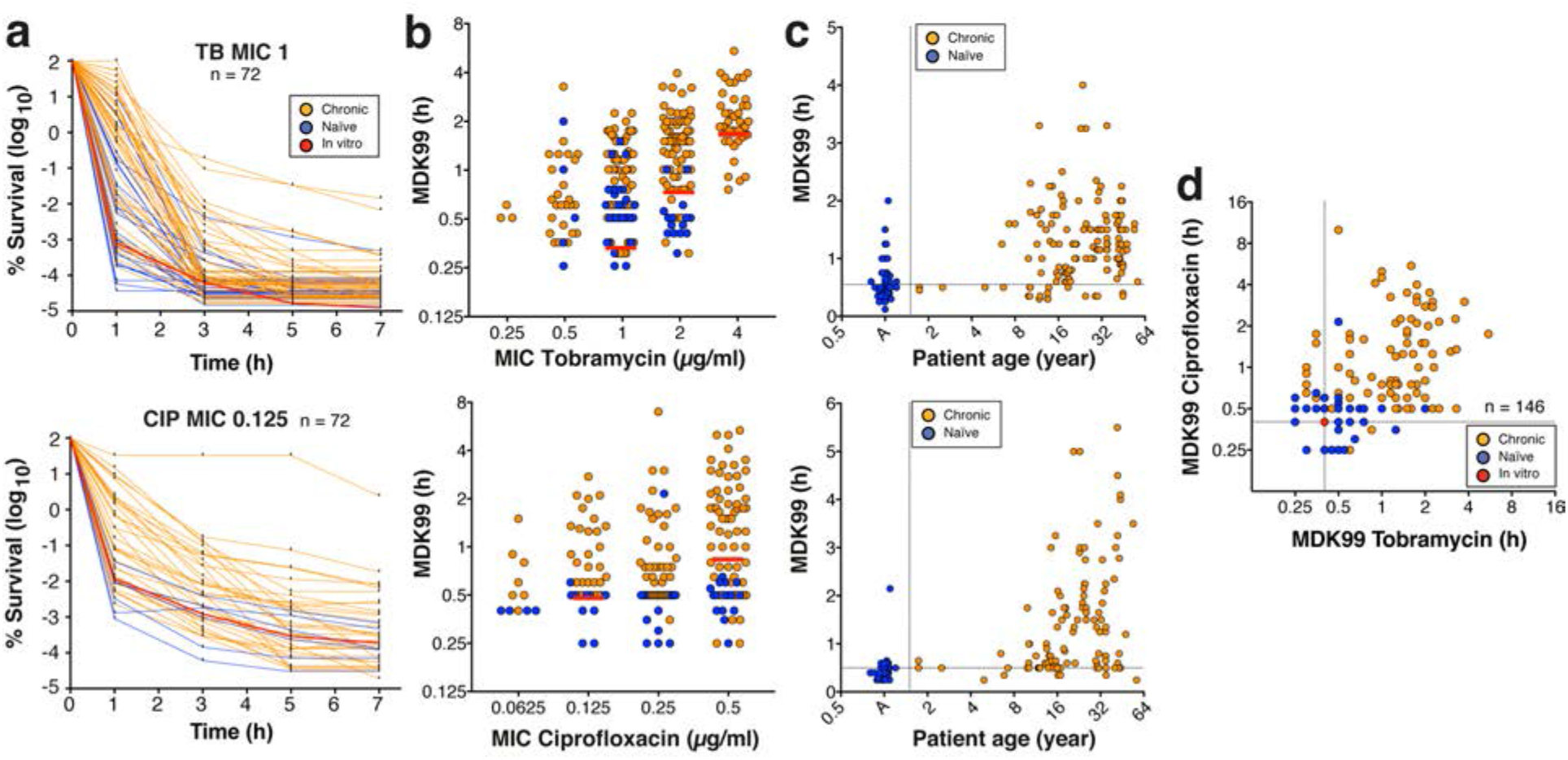
Selection of multi-drug tolerance during chronic infections in CF patients. *P. aeruginosa* isolates from acute (blue, n= 30) or chronic infections (orange, n= 81) were tested for survival when challenged with 32 μg/ml of tobramycin or 5 μg/ml ciprofloxacin. **a**) Killing kinetics of clinical isolates grouped according to their resistance levels (MIC). Survival rate is plotted over time. The red lines represent mean survival (± s.d., n= 6) of the control lab strain PAO1. Data were normalized to CFU at *t* = 0 hours. **b**) MDK99 values are plotted against the minimal inhibitory concentrations (MIC) of the respective isolates. MDK99 of *P. aeruginosa* laboratory strains with defined MICs are shown as red lines (mean ± s.d.). **c**) MDK99 values are plotted against the age of CF patients at the time of strain isolation. Isolates from acute infections (A) are shown in the first column. In a-c, upper and lower panels indicate tobramycin and ciprofloxacin treatment, respectively. **d**) MDK99 values of *P. aeruginosa* CF isolates treated with tobramycin and ciprofloxacin. Only isolates that were sensitive to both tobramycin (MIC <4) and ciprofloxacin (MIC <1) were considered (n =146). Survival of the *P. aeruginosa* lab strain PAO1 is indicated by a red circle. In a-d, the data are from 2 independent experiments.

Cluster analysis of the curve killing profiles delineated four different categories that were highly similar for both drug classes (Extended data Fig. 1b,c). Isolates from acute infections and control strains generally clustered in the first group including the most rapidly killed isolates and with the lowest levels of persisters. Also, CF isolates found in this group were from patients of the lowest age class (Extended Fig. 1c,d). In contrast, most isolates from chronic CF infections grouped to clusters 2-4, which are characterized by minor changes in MIC but largely increased MDK99 values and/or increased levels of persisters. Tolerance against tobramycin or ciprofloxacin was particularly pronounced in isolates from older patients (Fig. 1c).

From these data we concluded that the ability to survive antibiotic exposure is wide-spread in patient isolates and that increased tolerance may evolve during chronic infections, allowing pathogens to survive a broad spectrum of antibiotics during chemotherapy ^17,18^. The observed large differences in drug tolerance at any discrete level of antibiotic resistance re-iterates that resistance and tolerance, although contributing to the same resilience phenomenon, are distinct phenotypes.

### Sequential treatment with a single antibiotic selects for tolerance and resistance

The above data raised the question, which selective conditions would favor the evolution of tolerance over resistance development in *P. aeruginosa*. To explore the selective regimens leading to the evolution of antibiotic tolerance and resistance, *P. aeruginosa* cultures were treated at daily intervals with high levels of tobramycin (32 μg/ml = 32x MIC), concentrations that are readily observed in the sputum of CF patients during aerosolized drug treatment ^31^. Daily cycles included resuspension of parallel overnight cultures in fresh medium containing tobramycin, drug exposure for 3 hours, followed by washing and overnight growth in drug-free fresh medium (Fig. 2a). Bacterial survival was analyzed by plating after each killing phase, while MIC measurements and genome sequencing were performed each day after re-growth. Whereas less than 0.0001% of cells survived at the beginning of the experiment, all lineages rapidly adapted to the antibiotic regimen with a step-wise increase in survival rate. On average, tobramycin had lost its efficacy after 7 to 8 cycles, when all 13 independently evolved lineages had reached 10- to 32-fold higher MICs than the ancestor (Fig. 2b and Fig. 2c; Extended Data Fig. 2).

**Figure 2:**
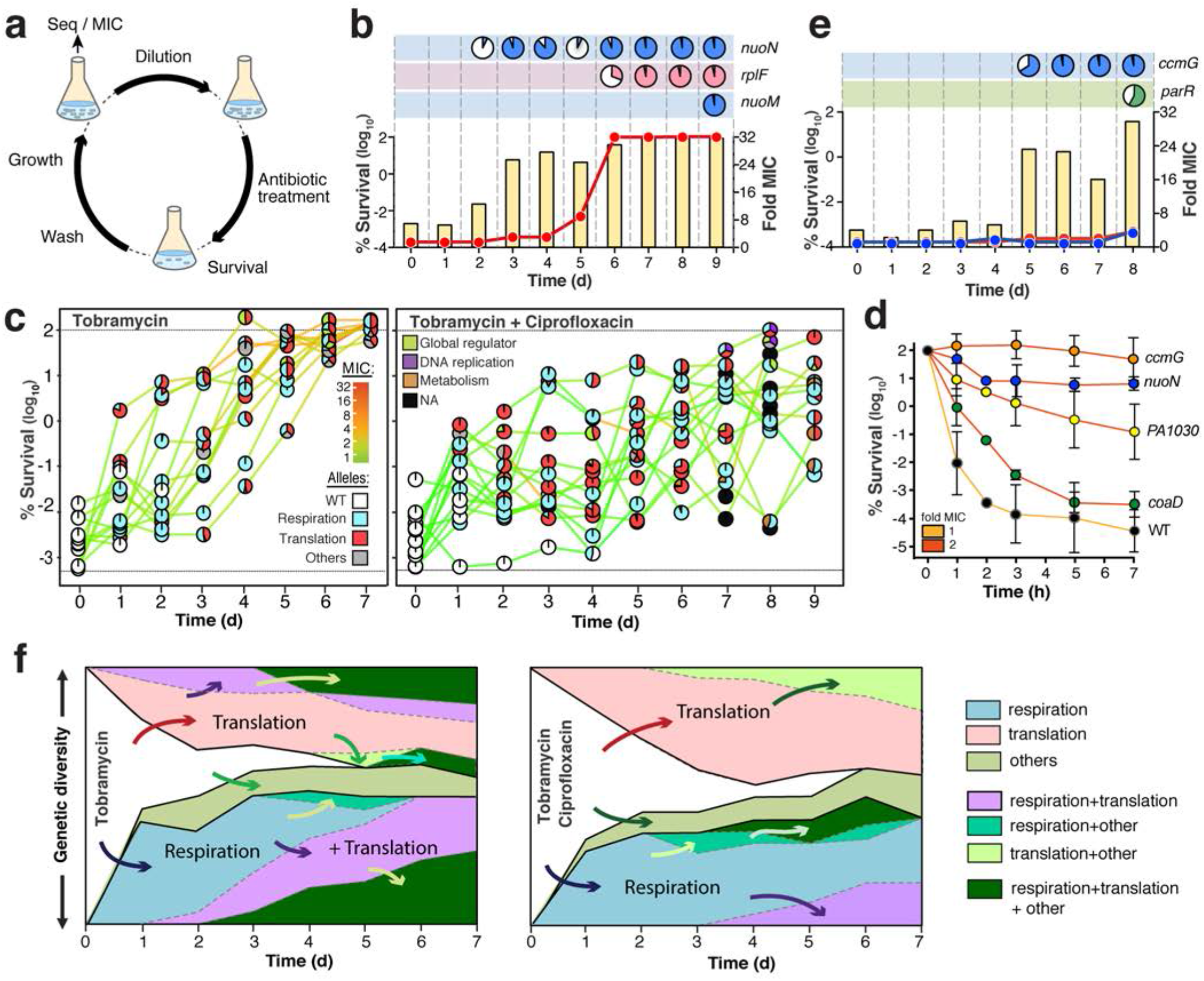
Evolution of antibiotic tolerance in *P. aeruginosa*. **a)** Experimental design for iterative exposure of *P. aeruginosa* to bactericidal antibiotics (for details see Materials & Methods). **b)** Example of *P. aeruginosa* response to cyclic exposure to tobramycin. The fraction of surviving cells (yellow bars) and MIC population values are indicated for tobramycin (red). Mutations acquired during selection are indicated in the boxes above the graph (see also: Extended Table 1; Extended Data Figs. 2, 5) with pie-charts representing population distribution of mutations (filled-area). **c**) Evolution of the survival of lineages treated with tobramycin (left) or with tobramycin/ciprofloxacin (right). Survival is plotted together with MIC values for tobramycin (colored lines). MIC values for ciprofloxacin can be found in Extended data Fig. 4. Acquired mutations are indicated by pie-charts with white circles indicating wild-type genomes and colors of invading alleles indicating specific functional classification. The lower dotted black line represents the detection limit for survival. **d)** Increased survival to tobramycin of selected tolerance mutants. Individual alleles were crossed back from evolved lineages (see Extended data Fig. 2) into the ancestral strain and survival of purified mutants was scored over time during treatment with tobramycin (32 μg/ml). The MIC is indicated by colored lines. Values are mean ± s.d. of at least n = 3 experiments. **e)** Example of *P. aeruginosa* response to cyclic exposure of tobramycin and ciprofloxacin. Details are as indicated in (**b**) with MIC values of evolving populations shown in red (tobramycin) and blue (ciprofloxacin), respectively. **f**) Evolutionary trajectories observed during selection with tobramycin (left panel) or with tobramycin and ciprofloxacin (right panel). The vertical axis indicates the proportion of lineages harboring mutations in genes of the indicated functional categories (genetic diversity). Independent evolutionary routes are separated by thicker black lines with successive mutational events being indicated by arrows. Functional categories of mutated genes are indicated on the right.

Time resolved genome sequencing uncovered the appearance of specific mutant alleles during tobramycin selection. Allele linkage was inferred from quantification of sequencing coverage of the respective alleles and was experimentally confirmed by PCR and targeted sequencing (Fig. 2c; Extended Data Fig. 2; Extended Table 1). The majority of the genetic changes were non-synonymous single nucleotide polymorphisms (SNPs) (Extended Table 1), implying a functional role of the respective proteins in antibiotic survival. Most mutations mapped to genes of few functional categories, including respiration and energy metabolism (*e.g. nuoN, nuoM, ccoN1, ccmF*), protein synthesis (*e.g. fusA* and *rplF*) and global regulation (*e.g. parRS* and *pmrAB*) (Extended Data Fig. 2, Extended Table 1). While some of these genes have been implicated in antibiotic tolerance ^9^, most of them are linked to target-specific ^32,31^ or indirect resistance mechanisms ^32,33^. In line with this, survival rates and MIC values increased in parallel in most lineages (Extended Data Fig. 2). From this and from the observation that, throughout most of the selection, MICs remained significantly below the concentration of tobramycin used in the experiment, we concluded that intermediate resistance levels boost population survival even at peak antibiotic concentrations. To test this, we isolated spontaneous resistance mutants of *P. aeruginosa* able to grow on agar plates with defined concentrations of tobramycin. Mutants with MICs of 1, 2, 4, and 8 μg/ml indeed showed gradually increasing survival during treatment with 32 μg/ml tobramycin (Extended Fig. 3a). These findings underscore that low-level resistance below the clinical resistance breakpoint can strongly promote survival at high therapeutic doses, and emphasize that antibiotic tolerance should always be assessed in a resistance-neutral context, *i.e.* by comparing strains with similar MICs (Fig. 1a,b). In addition, resistance- and tolerance-mediated survival benefits can be distinguished based on their drug-specific or multi-drug phenotypes, respectively.

Genome sequencing revealed the sequential invasion and fixation of specific alleles during daily treatment intervals (Fig. 2b,c) with SNPs in genes involved in respiration and energy metabolism generally preceding the acquisition of resistance mutations (Extended Data Fig. 2). To test their role in survival, a selection of the respiration-related alleles was crossed back into the ancestral background. While some alleles showed moderate effects (e.g. *nuoD*, *ccmF*) (Extended Data Fig. 3b), a point mutation in *nuoN* (G300D) encoding a subunit of the respiratory complex I strongly increased survival during tobramycin (Fig. 2d) and during ciprofloxacin treatment (Extended Data Fig. 3c). The *nuoN* mutant showed an extended lag-phase (on average 7h long) when exiting from stationary phase, a phenomenon typically observed in hyper-tolerant strains (Extended Data Fig. 3d) ^9,10^. Remarkably, the tolerance phenotype of this lineage was lost by a second site mutation in *nuoM* after penetration of a mutation in *rplF* conferring high-level resistance to tobramycin (Extended Fig. 2, Lineage 1; Extended Fig. 3e). Thus, tolerance mutations can impose fitness costs, which can be compensated through second site mutations after populations have reached high levels of resistance. It is possible that tolerant variants transiently invade pathogen populations during antibiotic therapy, explaining the difficulties of gauging the actual contribution of tolerance to resistance development and treatment failures.

In sum, these results indicated that, in contrast to what had been reported for *E. coli ^20^*, single drug treatment drives parallel evolution of tolerance and resistance in *P. aeruginosa*.

### Sequential treatment with a drug combination strongly selects for tolerance

We next challenged *P. aeruginosa* with a combination of tobramycin and ciprofloxacin at therapeutic doses. Survival again rapidly increased in multiple independent lineages with killing efficiencies dropping four to five orders of magnitude after 7-9 cycles. In contrast to single drug treatment, all lineages evolved high levels of tolerance against both tobramycin and ciprofloxacin without developing significant levels of resistance (Fig. 2c,e; Extended Data Fig. 4; Extended Data Fig. 5a,b; Extended Table 1). Tolerant lineages often developed sub-populations with extended lag phase or slower growth (Extended Data Fig. 5c). In contrast to most lineages obtained during single drug treatment, double-drug treatment generated lag phases (on average 3h) that precisely matched the duration of drug exposure, indicating evolutionary trajectories minimizing fitness costs under these conditions. Several of the tolerant lineages showed cross-tolerance against membrane active compounds like polymyxin B (Extended Data Fig. 5d) and this phenomenon was also observed in lineages evolved with tobramycin alone. Thus, treatment with a combination of bactericidal antibiotics strongly favors the development of multidrug tolerant *P. aeruginosa* ^2^. The observation that some of the tolerant lineages acquired intermediate-level resistance against both drugs at later time points (Extended Data Fig. 4, lineage 16, 17, 20, 23, 25), indicated that during combination therapy, tolerance-based survival may promote the development of multidrug resistance ^11,33^.

Time resolved population sequencing and PCR-based SNP confirmation of individual clonal members, showed that sub-lineages co-existed for several selection cycles and generally conferred a large increase in survival (Fig. 2e; Extended data Fig. 4; Extended Table 1). Crossing back specific alleles into the ancestral lineage revealed their multi-drug tolerance phenotype as indicated by large increases of survival (Fig. 2d; Extended data Fig. 3c and 5a,b). Most mutations that increased tolerance against both drugs led to a small increase of resistance to tobramycin (MIC 2 μg/ml^−1^) but not ciprofloxacin, an effect that cannot account *per se* for the large increase in tolerance (see MIC controls, Extended data Figs. 3a and 5a). Also, several tolerant strains showed reduced MIC for ciprofloxacin (Extended data Fig. 5e), suggesting that under these conditions, tolerance development is strongly favored. Similar to the single drug regimen, tolerance alleles mapped to genes involved in respiratory and energy metabolism, including respiratory complex I, cytochrome c maturation and the cytochrome c oxidase complex (Fig. 2c,d; Extended Table 1). In line with this, altered respiration had recently been linked to increased persister levels in *E. coli* ^34^. Finally, SNPs in genes coding for a toxin-antitoxin system (*PA1029*-*PA1030*) and cell metabolism (*coaD)* were associated with increased tolerance. *PA1030* encodes a toxin of the RES domain family that was recently shown to interfere with NAD metabolism ^35–37^.

Typically, tolerant mutants evolved directly from a susceptible ancestor and were able to outcompete other co-emerging linages, irrespective of differences in MIC (Fig. 2c; Extended Data Fig. 4, lineages 13, 19 and 22). Once established, such lineages provided the genetic background for the fixation of additional mutations that either conferred low level resistance (*e.g. gyrA* ^*38*^ in lineage 20; *rplF* ^39^ or *fusA* in lineage 22; *parR* ^*40*^ in lineage 25) or further increased tolerance (*e.g. nuoN* in lineage 15, or *hxcR* in lineage 19) (Extended Data Fig. 4). Alleles conferring substantial resistance against one drug could invade transiently, but were not stably maintained under these conditions (*e.g.* lineages 14, 17). Several SNPs mapped to *fusA1*, the gene for elongation factor G (EF-G) (Extended Data Figs. 4, 5e; Extended Table 1), a protein that was recently associated with both tolerance and aminoglycoside resistance *in vivo* ^32,41,42^. Most *fusA1* alleles conferred only moderate increases of survival in both tobramycin and ciprofloxacin and were often outcompeted by other sub-lineages (Extended Data Fig. 4, 5e).

In sum, these experiments revealed that although selection with one or two antibiotics yielded different mutations, similar genes and functional pathways were affected under both conditions. In general, the acquisition of mutations in components of the translation apparatus was preceded by mutations in genes involved in respiratory process or vice versa (Fig. 2e). These results suggested that these two pathways synergistically influence antimicrobial sensitivity and survival, possibly via their central role in the generation and consumption of energy.

### Respiratory mutations generate slow-growing subpopulations

To better understand antibiotic tolerance, a selection of the mutants emerging from experimental evolution was characterized in more detail. *E.g.* a Gly to Asp exchange in the NuoN transmembrane subunit of the Respiratory Complex I (RCI) conferred high levels of tolerance and increased levels of persisters (Fig. 2d, Extended data Fig. 5e). This Gly residue is positioned in the immediate vicinity of helix 7 of NuoN, a region known to be involved in proton-pumping activity ^43^. In line with this, we found that a subpopulation of *nuoN* mutants had reduced membrane potential (Fig. 3a). Because up-take of aminoglycosides depends on the membrane-potential ^44^ this could explain tobramycin tolerance but not tolerance against ciprofloxacin which up-take is independent of the proton motive force^45^. Alternatively, cells with low membrane potential may have reduced growth rates or extended lag phases (Extended fig. 3d) during outgrowth in fresh medium. In line with this idea, we observed distinct subpopulations of slow growing bacteria when analyzing the *nuoN* mutant expressing TIMER^bac 46^ as an indicator for growth rates (Fig. 3b). Of note, mutations in *nuo* genes were found in several lineages of *P. aeruginosa* isolated from CF patients, indicating that these genes are under selection *in vivo* ^47^. Future work will need to correlate drug tolerance with slow growth and/or extended lag phases of these mutant strains at the single cell level.

**Figure 3:**
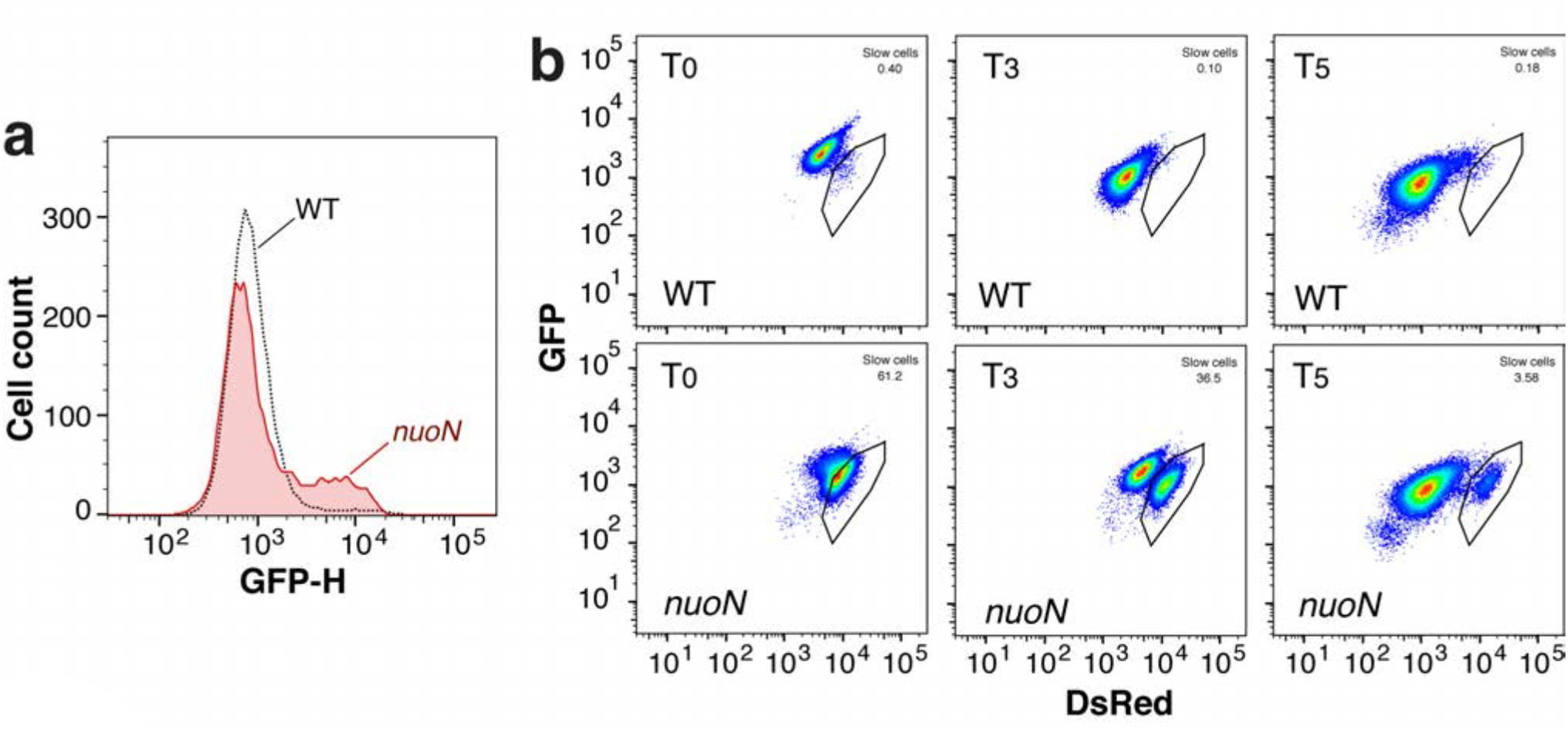
The *nuoN* hyper-tolerant mutant strain shows a subpopulation with low membrane potential and reduced growth rates. **a**) Flow cytometry analysis of overnight cultures of PAO1 wild type and isogenic *nuoN G300D* mutant incubated with DiBAC4, an indicator dye of membrane potential that is only taken up by cells with collapsed membrane potential ^59^. **b**) Flow cytometry of PAO1 wild type and isogenic *nuoN G300D* mutant containing a plasmid with constitutive expression of TIMER^bac 46^. Bacteria were grown overnight, diluted into fresh medium and analyzed by FACS over time. Recordings are shown at time point 0 and after 3 and 5 hrs of incubation, respectively. Black outlines indicate slow-growing sub-populations that were observed only in the tolerant mutant strains.

### Tolerance facilitates resistance development in *P. aeruginosa*

The above results indicated that tolerance evolves rapidly when *P. aeruginosa* is periodically exposed to high concentrations of antibiotics and that tolerance development often precedes resistance. To test if tolerance influences the rate of resistance development, we periodically exposed *P. aeruginosa* wild type and mutants with distinct levels of tolerance (low, medium, high) to tobramycin (Extended data Fig. 6a,b). While all strains gradually evolved resistance over time, the rate of resistance acquisition was similar in all strains (Fig. 4a; Extended Data Fig. 6a; Extended Table 1). However, tolerant lineages were more likely to survive the initial selection as compared to the non-tolerant PAO1 lab strain (Fig. 4b), indicating that tolerance provides bacterial populations with a significant advantage to evolve resistance. To test this, we performed antibiotic selection experiments with a 1:1 mixture of PAO1 wild type and isogenic high-tolerance variants (Fig. 4c). We found that PAO1 was consistently outcompeted by its isogenic high-tolerance counterpart when mixtures were repeatedly exposed to high concentrations of tobramycin. Resistance was systematically acquired by strains exhibiting the highest level of tolerance, even when co-evolved with non-tolerant strains with higher initial MIC levels. Even in the few rare cases where the original low-tolerance competitor strain was still detected at the end of the treatment regimen, mutations conferring resistance were acquired by the high-tolerance ancestor (Fig. 4c). Importantly, while the hyper-tolerant *nuoN* mutant was highly successful in acquiring resistance mutations, its low-tolerance isogenic descendant *nuoN nuoM* (Extended data Fig. 3e) failed to acquire resistance (Fig. 4c).

**Figure 4:**
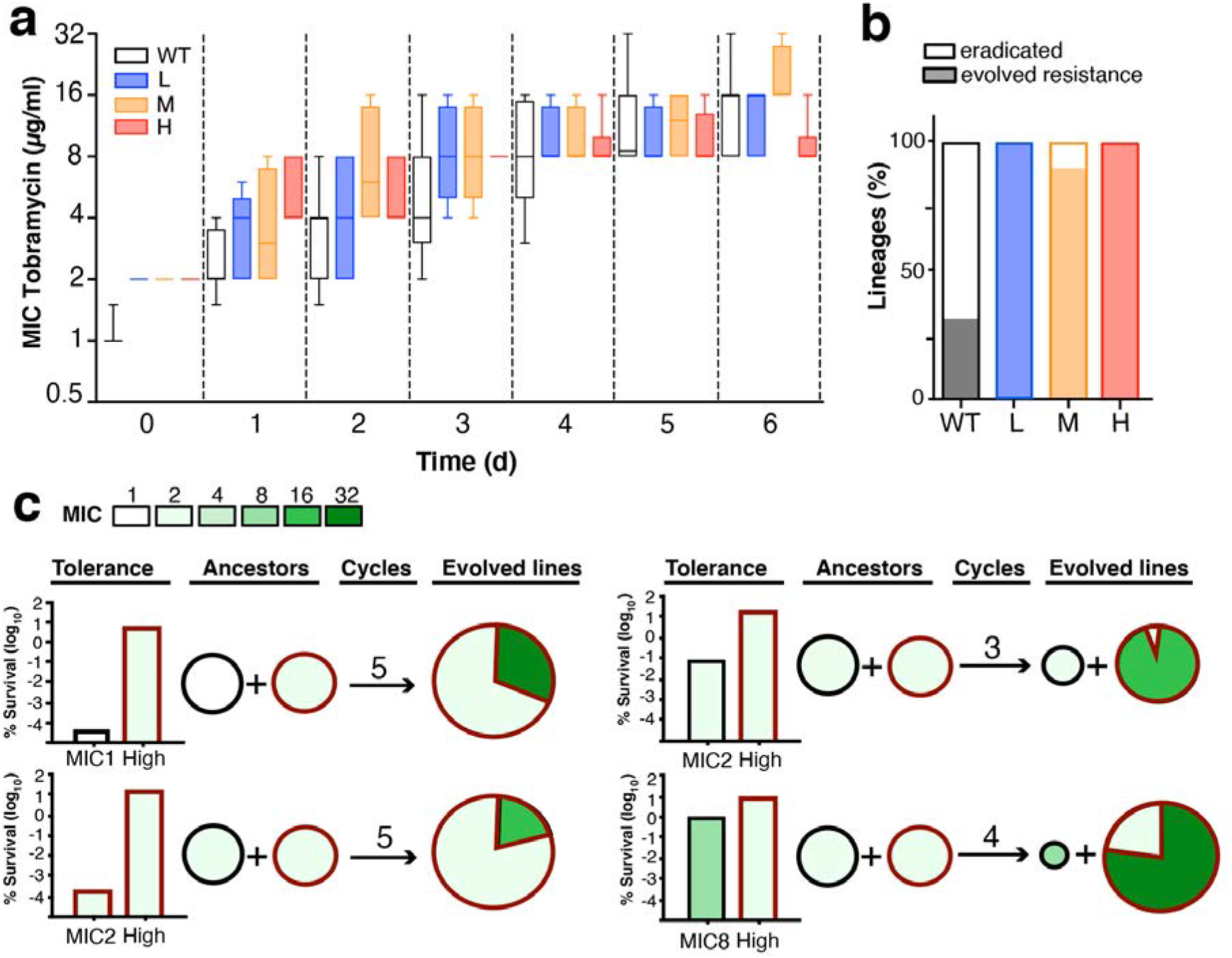
Tolerance facilitates antibiotic resistance development. **a)** Tolerance does not accelerate resistance development. Resistance (MIC) was evolved over time in strains with different initial levels of tolerance (H=high , *nuoN*_*G300D*_; M=medium , PA1030*; L=low , *nuoD*_*R551P*_; see: extended data Fig. 5) by serial daily exposure to tobramycin (32 μg/ml). Box plots are shown with quartiles and standard deviation of at least 3 experiments. **b)** Tolerance facilitates the evolution of resistance lineages. Strains with different levels of tolerance as in a) were analyzed during daily exposure to tobramycin. The fraction of independent lineages that were eradicated (white) or developed resistance (black) during drug treatment are indicated (n=6). **c)** Tolerance provides a strong advantage during development of antibiotic resistance. 1:1 mixtures of low (black) and high tolerance (red) *P. aeruginosa* strains were challenged daily with tobramycin (32 μg/ml). Tolerance levels of the initial strains are indicated as survival after 3h of drug treatment (left). Daily treatment intervals were as described in Fig. 2a with the number of selection cycles indicated for each experiment. Genetic changes were determined by whole population genome sequencing at the end of the selection. Resistance levels and resistance allele distributions are indicated. Red and black frames indicate resistance evolution in the high-and low-tolerance ancestors with initial MIC levels indicated.

From this we concluded that tolerance provides *P. aeruginosa* with a strong selective advantage when challenged with bactericidal antibiotics by promoting survival and by increasing the probability of resistance development. Tolerance may not only increase the chance of resistance alleles to emerge and spread, but may also increase the stringency of selection as emerging resistant variants need to be able to outcompete higher numbers of persisting individuals during periods of unconstrained growth.

### Evolution of tolerance and resistance during chronic infections of *P. aeruginosa*

To better understand the evolutionary trajectories leading to antibiotic resilience in human patients, we reconsidered the resistance and tolerance of 252 drug sensitive isolates of *P. aeruginosa* derived from a cohort of 91 CF patients. To be able to assess resistance and tolerance in parallel, we analyzed only strains that had retained sensitivity to at least one of the two drugs used in this study. We determined resistance (MIC), tolerance and persistence for individual patient isolates or from patients sampled repeatedly over periods of up to ten years. Visualizing these phenotypic distributions across patient age at the time of isolation illustrates the evolutionary trends of antibiotic resilience in the clinical dataset. In order to avoid resistance effects in the assessment of tolerance and persistence, we reported resistance against an antibiotic A vs. tolerance or persistence against an antibiotic B and *vice-versa* (496 comparisons; Fig. 5a).

**Figure 5:**
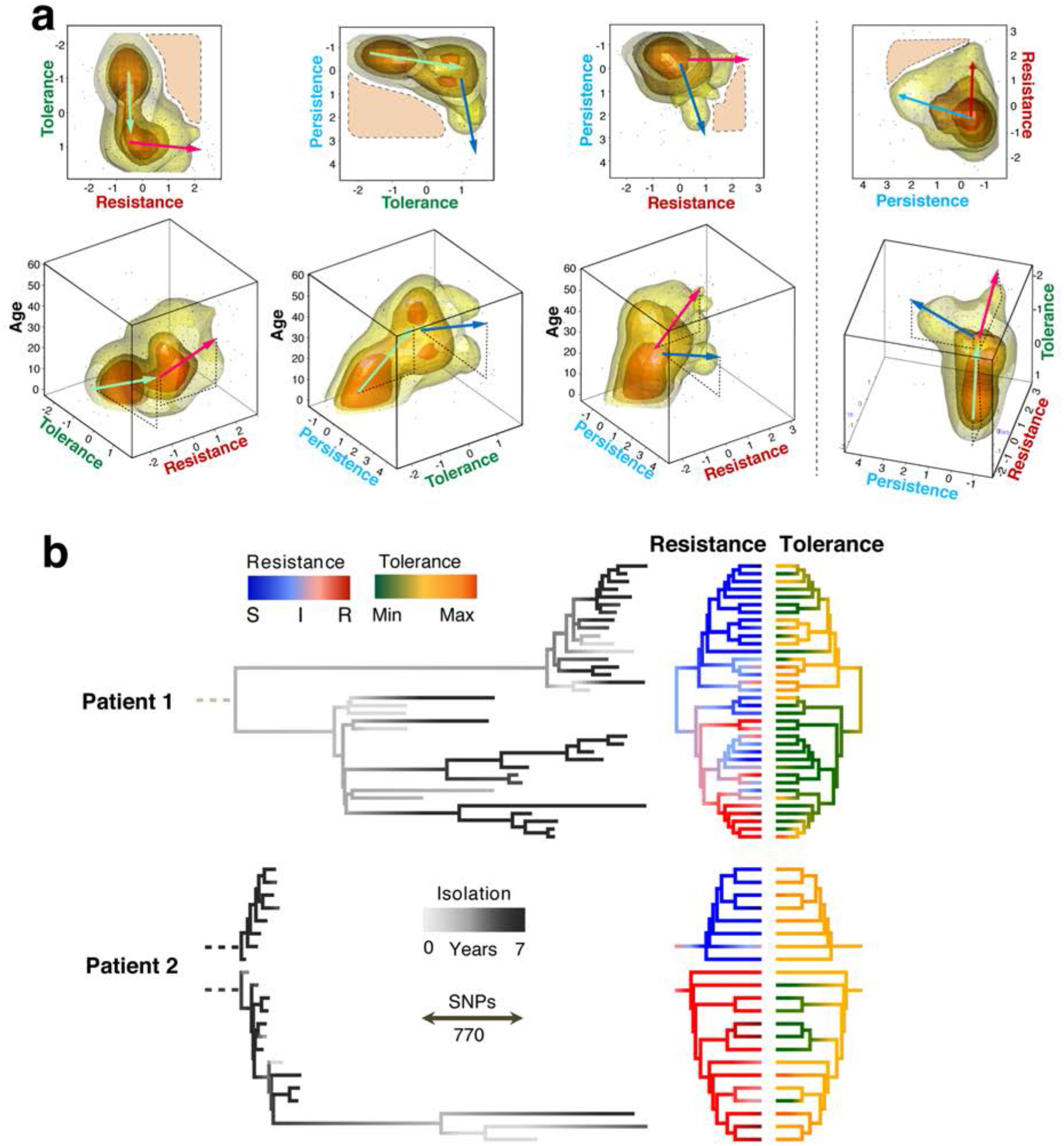
Microevolution of tolerance, persistence and resistance during chronic *P. aeruginosa* infections. **a**) Illustration of antimicrobial phenotypes during host microevolution. Strains (dots) and their density distribution (clouds) are represented as function of three antimicrobial parameters and patient age at the time of strains isolation: tolerance (survival at 1h treatment), persistence (survival at 7h treatment) and resistance (MIC). Naïve (n=67, age = zero) and chronic isolates of *P. aeruginosa* are considered (n=312, 91 patients from five to 61 years old) (see also Extended Data Figure 7). In order to eliminate resistance effects on tolerance and persistence, we report resistance against an antibiotic A vs. tolerance or persistence against an antibiotic B. Only strains sensitive to drug B (tobramycin: MIC<4μg/ml or ciprofloxacin: MIC<1μg/ml, n=496 comparisons) are reported here. Tolerance and persistence are shown as log10 survival values standardized for drug B. Resistance values correspond to Log2(MICs). Cloud colors indicate data densities with numbers increasing from yellow to orange. Arrows indicate phenotype progression with color indicating with different parameters (green = tolerance; blue = persistence; red = resistance; black = age of patient). Areas with dashed outlines delineate phenotype profiles that were rarely observed. **b)** Genome-based phylogeny (85.579 variant positions) of *P. aeruginosa* strains from two CF patients. The branch lengths of the trees on the left indicate genetic differences (SNPs). Dashed roots represent the extrapolated clonal ancestors. Colors of the terminal branches indicate isolation time within a five years window (grey scale); resistance (sensitive, blue; intermediate, white; resistant, red); and tolerance (lowest tolerance, green; intermediate tolerance, yellow; highest tolerance, orange). Resistance is plotted as the average log2 fold variation from intermediate MICs for five antibiotics (Extended Data Figure 7). Tolerance is plotted as the average survival scores for antibiotics, for which no resistance was observed.

Strains isolated from younger CF patients generally displayed low resistance and low tolerance. With increasing age, the number of drug tolerant isolates gradually rose while resistance to tobramycin or ciprofloxacin remained low (Fig. 5a). Hyper-tolerant but susceptible (low MIC) strains were often found in patients at intermediary age (15 to 30 years old) while increased numbers of resistant isolates were only observed at higher patient age (above 30 years old) and generally lagged behind tolerance development (Fig. 5a). Intriguingly, we observed an accumulation of two distinct pathogen subpopulations in older patients: isolates that were highly resistant to both antibiotics and isolates that had retained low level resistance but had acquired a hyper-persister phenotype (Fig. 5a). These observations indicated that *P. aeruginosa* gradually adapts during chronic infections by first evolving increased antibiotic tolerance, followed by an increase of multidrug resistance or hyper-persistence at later stages of the infection. Importantly, while moderate levels of persistence and resistance coexisted in some isolates, strains displaying both high levels of persistence and resistance were rarely observed (Fig. 5a). In fact, the distribution of persistence and resistance in clinical isolates from late stages of infection as depicted in Fig. 5a (top right plot) suggests that both phenotypes contribute to the same selective phenomenon.

Next we investigated resistance and tolerance phenotypes of *P. aeruginosa* isolates collected longitudinally from two CF patients over several years (Fig. 5b; Extended Data Fig. 7). Whole genome sequences of 58 isolates were compared to references from GenBank revealing highest similarities with strain RIVM-EMC2982, a clinical isolate from the Netherlands (CP016955.1), 12-4-4(59), a strain isolated from blood of a burn patient (NZ_CP013696.1 ^48^) and W36662, a strain isolated from a cancer patient (CP008870.2). All strains from patient 1 belong to a single clonal clade (W36662), while sequences of isolates of patient 2 indicated a recent and ongoing superinfection by different clones (RIVM-EMC2982 and more recently 12-4-4(59)). In line with previous observations ^13^, parallel evolution of coexisting *P. aeruginosa* lineages was observed in both patients. As expected, longitudinal sampling did not directly reflect genetic evolution as more recent isolates did not necessarily descended from previous isolates but rather emerged from a common ancestor ^13^. Phylogenetic analyses indicated that in patient 1, an early split of the clonal lineages separated populations into two clades with similar distribution during sampling over the next years (Fig. 5b). Given the significant evolutionary distance between these clades, we cannot distinguish whether they are the result of microevolution of a common ancestor or a coinfection by two closely related isolates.

In both patients, the highest levels of resistance against different classes of antibiotics (tobramycin, ciprofloxacin and meropenem) were observed within the same phylogenetic branch, while the other lineages showed much lower levels of resistance but significant levels of tolerance (Fig. 5b). Isolates with low MIC but high-tolerance showed similar survival as resistant isolates, underscoring that tolerance is highly successful during chronic infections. In both patients, the genetic divergence (*i.e.* branch length) was significantly higher in the clades exhibiting high-level resistance irrespective of isolation time (Fig. 5b), suggesting that different resilience phenotypes may result from different levels of genetic alterations. This is in line with the multidrug nature of tolerance (Fig. 1d), which may represent a simpler genetic solution during multi-drug therapy as compared to evolving multiple drug-specific resistance mechanisms. Importantly, multidrug resistant strains that retained sensitivity to one specific drug (e.g. tobramycin), generally displayed high-level tolerance against this antibiotic (Extended Data Fig. 7).

Although the sample density did not permit identifying genetic variations correlating with antibiotic tolerance or resistance by genome-wide association, sequential acquisition of known resistance alleles (e.g. *fusA1*, *pmrB*, *gyrA*, *gyrB*, *oprD*) ^49^ or mutations in genes that were shown in this study to enhance tolerance (*nuoG*, *nuoN*, *ccmG*, *PA1030*, *PA1549*) were readily observed in the clinical isolates (Extended Data Fig. 8). Together, these phylogenetic analyses propose that different antimicrobial survival strategies (i.e. tolerance/persistence and resistance) evolve and exist in chronically infected patients. Our data also argue that at later stages of chronic infections of CF airways, mutually exclusive evolutionary trajectories result in increased resistance and persistence, respectively.

## Discussion

Based on our findings, we propose a dynamic evolutionary model of acquisition of tolerance and resistance during antibiotic chemotherapy. Typically, chronic infections are sparked by a naïve non-tolerant and sensitive strain. Despite of early treatment with antibiotics, initial colonization of CF airways does not seem to result in resistance development in *P. aeruginosa* populations ^50^. Our data argue that antibiotic exposure selects for increased tolerance early during CF lung colonization in particular during treatment with combinations of antibiotics administered jointly or serially ^11,51^. Preferential development of tolerance over resistance may be due to a wide range of possible tolerance mechanisms, providing a larger target size for mutations alleviating antibiotic stress ^9^. Moreover, due to its multi-drug phenotype, tolerance development may be particularly favored during chemotherapy with combinations of drugs, a treatment strategy that is often used for CF patients by combining inhalation and systemic therapy ^29^. In agreement with this, we show here that recurrent exposure of *P. aeruginosa* to high doses of tobramycin and ciprofloxacin leads to the rapid evolution of hyper-tolerance but not high-level resistance. In contrast, resistance is generally drug specific with mutations being commonly limited to direct drug targets. Once established, tolerance likely provides populations with a significant survival advantage, facilitating the acquisition of genetic changes that further increase antibiotic resilience ^11,20,52^. Our studies indicate that during chronic infections in CF lungs, this process ultimately leads to genetic changes conferring high-level resistance against multiple antibiotics or to further increments of tolerance levels and persistence (Fig. 5a,b). Independent evolutionary trajectories leading to different mechanisms of antibiotic resilience are in line with the observation that isolates with strongly diverging resistance levels are observed in individual patients, a phenomenon that may relate to physical separation and local microevolution in CF lungs ^13,14^.

Although antimicrobial tolerance is underappreciated and still largely neglected clinically ^4^, we propose that it plays an important role during in-host microevolution leading to rapid pathogen adaptation and the discharge of antibiotic stress. Our data imply that tolerance can not only precede and promote resistance development during early stages of infections but that it can also serve as an alternative mechanism enabling long-term survival of pathogens during continued antibiotic exposure during chronic infections. The observation that some of the isolates from older CF patients had retained low MIC and low tolerance against different antibiotics, argues for additional protective mechanisms such as pathogen encapsulation into biofilm structures ^53^. Limited bacterial intermixing and different selective pressures in different lung regions may explain much of the differences in tolerance and resistance profiles of isolates from the same patient ^13^.

In addition to the fitness advantage conferred by antibiotic tolerance, low-level resistance below the clinical breakpoint of antibiotics may contribute to survival during chronic CF infections. Genetic studies indicated that the target size for mutations conferring low-level tobramycin resistance in *P. aeruginosa* is considerable ^54^. Moreover, mutations conferring low-level tobramycin or ciprofloxacin resistance also confer substantial fitness advantages at antibiotic concentrations in the sub-MIC range ^55^. In this study, we showed that low-level resistance also contributes to increased survival at very high antibiotic concentrations. Thus, mutations conferring low-level resistance show a tolerance phenotype at concentrations above their MIC. This argues that the contribution of resistance below the clinical breakpoint to pathogen survival should not be neglected as it may well help explain clinical treatment failures ^56^. It is also possible that the ‘tolerance aspect’ of low-level resistance may help explain their role in the development of high-level resistance ^57^. While we observed that mutations conferring tolerance often resulted in small increases of MIC, their effect on survival was disproportionally higher as compared to *bona fide* resistance mutations with similar MIC values. Moreover, they generally showed a multidrug phenotype, while mutations causing resistance were generally drug specific. Thus, despite of similarities in promoting survival during antibiotic treatment, tolerance and low-level resistance are clearly distinct phenomena and based on different molecular and cellular mechanisms.

Altogether, our findings shed new light on the forces shaping antibiotic resistance development in the human pathogen *P. aeruginosa* and emphasize the need for compounds with improved capacity to eliminate tolerant cells ^58^. Our results may also help optimizing antibiotic treatment strategies during chronic infections to minimize the risk of resistance development. For example, because tolerance influences the rate of *de novo* emergence of single and double resistances, detailed knowledge about tolerance development during chronic infections should influence treatment decisions regarding mono-drug vs. multi-drug therapy ^33^. Access to such information will require the development of simple and rapid tools for routine monitoring of antibiotic tolerance alongside with resistance.

## Acknowledgments

We thank M. Bläsi for assistance with generating data; K. Jahn (Pneumology, University Hospital Basel) for assistance with clinical isolates; R. Vesco, C. Kiessling, E. Schultheiss, M. Schneider, C. Straub, D. Wüthrich (Clinical Bacteriology and Mycology, University Hospital Basel) and C. Beisel (Genomics Facility Basel, ETH Zurich Department of Biosystems Science and Engineering) for generous help with the sequencing of evolved strains and clinical isolates. We are indebted to D. Bumann, C. Dehio, R. Neher and B. Laventie for valuable discussions and critical comments on the manuscript. This study was supported by the Swiss National Science Foundation NRP72 grant 407240_167080 to U. J. The authors declare no competing interests.

## Extended data figures

**Extended Data Figure 1:**
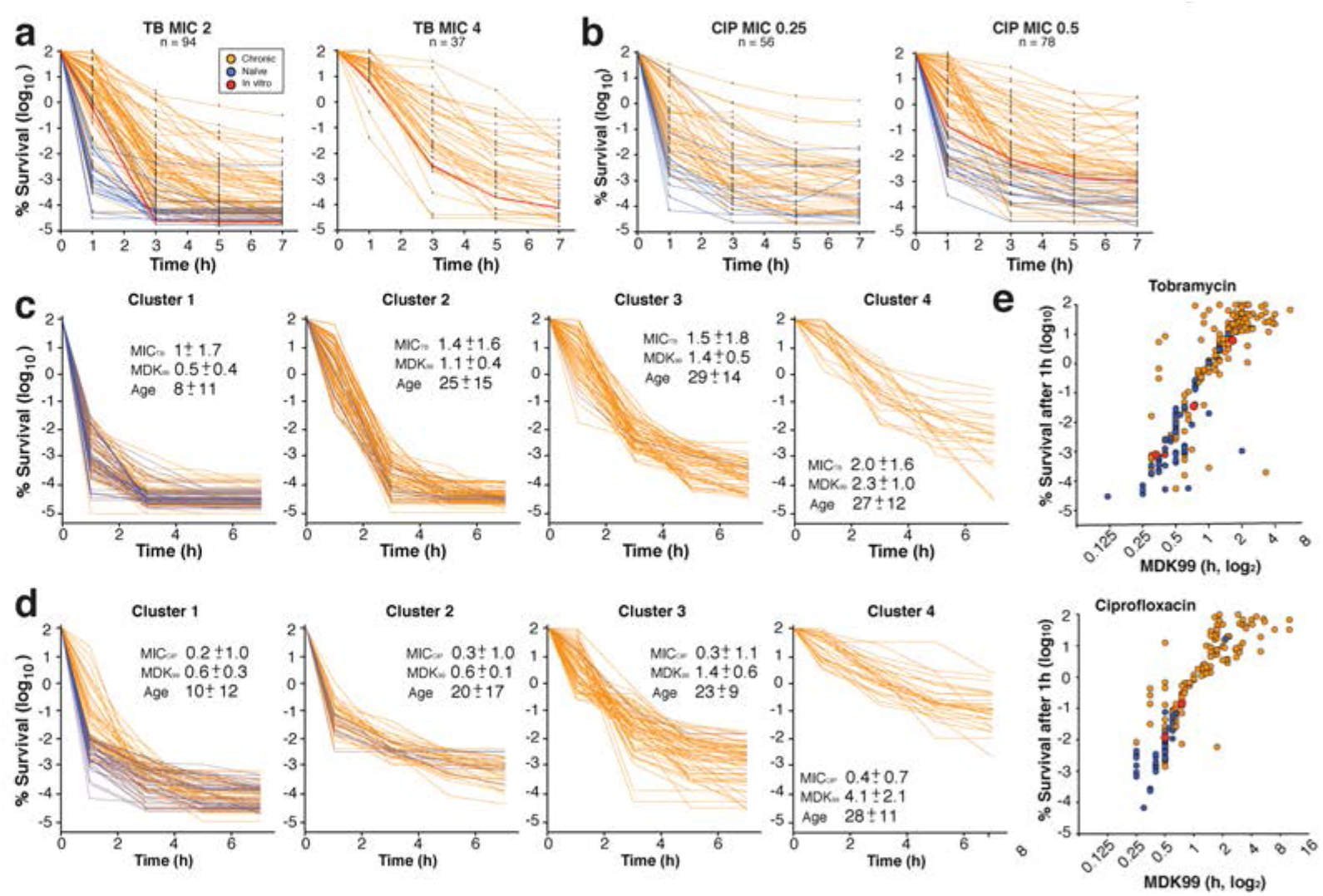
Increased tolerance and persistence in isolates from patients with chronic lung infections. *P. aeruginosa* acute infection isolates (blue, n= 58) or isolates from chronic infections (orange, n= 290) were tested for survival when challenged with 32 μg/ml of tobramycin or 5 μg/ml ciprofloxacin. **a)** and **b**) Killing kinetics of clinical isolates grouped according to their resistance levels (MIC) and treated with tobramycin (**a**) or ciprofloxacin (**b**). Survival rates are plotted over time. Red lines represent mean survival values (± s.d., n=6) for the *P. aeruginosa* lab strain PAO1 resistant variants with different MICs. Data were normalized to CFU at t = 0 hrs. **c, d**) Killing curves of *P. aeruginosa* isolates treated with tobramycin (**c**) or ciprofloxacin (**d**) were binned based on the decay of survival. Average MIC, MDK99 and sample age (± s.d.) are shown for each cluster. **e)** Survival of individual isolates after 1hr of killing with tobramycin (32 μg/m) or ciprofloxacin (5 μg/ml) plotted against MDK99.

**Extended Data Figure 2:**
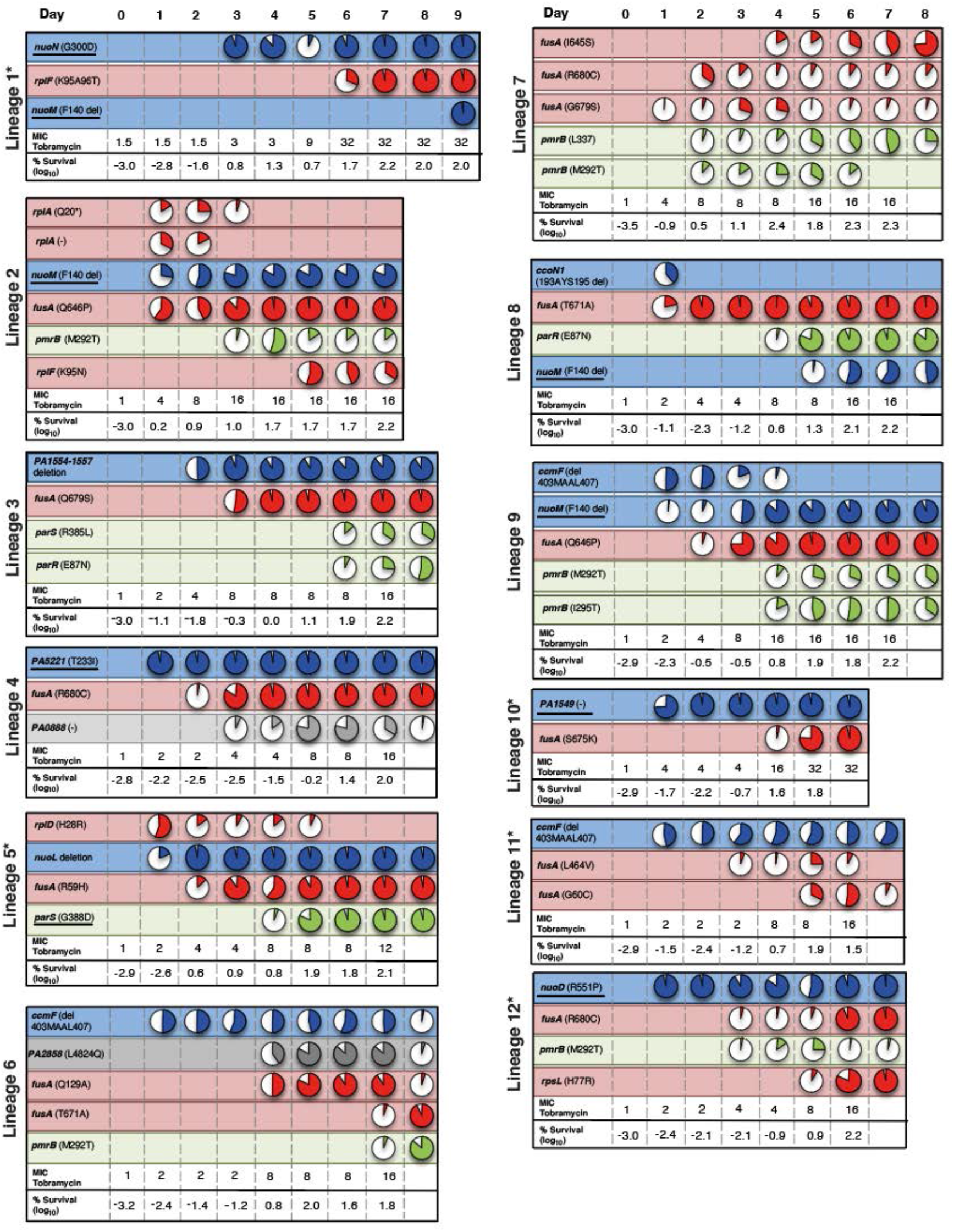
Mutations arising during single drug selection as identified by whole genome sequencing. Mutations in genes involved in respiration, protein synthesis, global regulators, DNA replication or other functions are indicated in blue, red, green, orange and grey, respectively. Circles indicate that a specific mutation was detected and filled areas in the pie-charts indicate mutation distribution in each lineage. For marked lineages (*) single colonies were randomly picked and allele combinations were analyzed by PCR. Specific mutations (underlined) were crossed back into the ancestral background and analyzed for their contribution to increased survival and MIC. MIC values and survival rates are reported for each day.

**Extended Data Figure 3:**
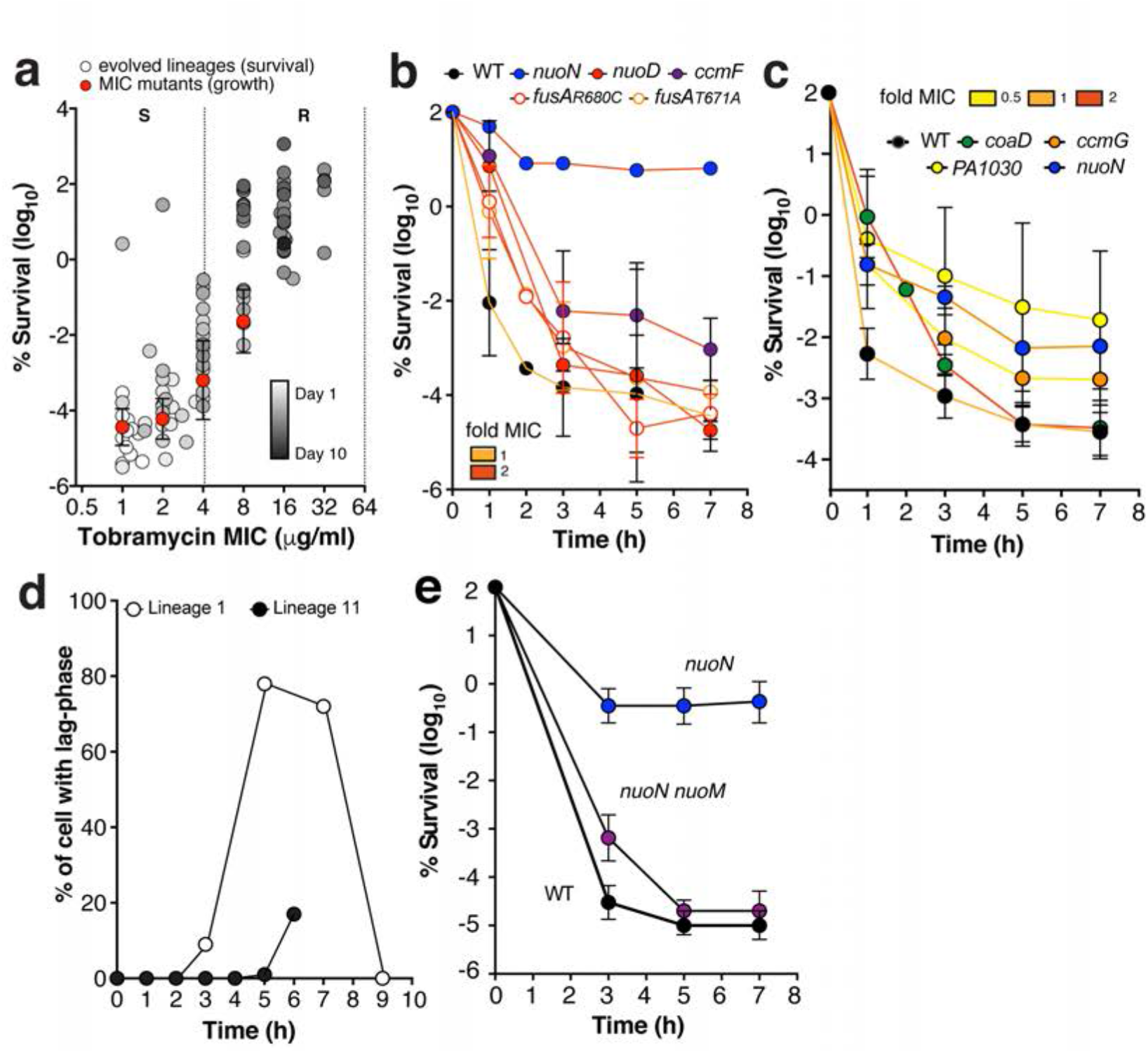
*P. aeruginosa* adaptation to tobramycin. **a)** Lineages evolved in the presence of tobramycin (see Extended data Fig. 2) were tested for tobramycin mediated killing (32 μg/ml). Survival after 3 hrs of treatment is plotted against MIC values of the evolved population (*n* = 90). The time of isolation (days) of the evolved strains is indicated by the grey scale. Survival of non-adapted lab strains with different MICs (*n*=12) is indicated by red dots (mean ± s.d.) (see Extended Data Fig. 1a). The dotted lines mark the breakpoints of resistance according to EUCAST^30^: S, sensitive; R, resistant. **b**) Contribution of individual mutant alleles to survival during antibiotic treatment. Tolerance alleles from strains isolated during selection (Extended Data Fig. 2) were crossed back into the ancestor and corresponding mutants treated with tobramycin (16 μg/ml). Survival is shown as mean ± s.d. of at least three experiments. Line coloring indicates MIC. Data in **a)** and **b)** were normalized to CFUs before treatment. **c**) Contribution of individual mutant alleles to survival during ciprofloxacin treatment. Selected alleles from evolved strains (Extended data Fig. 2) were crossed back into the ancestral strain and the corresponding mutants were treated with ciprofloxacin (2.5 μg/ml). Line coloring represents MIC values of individual strains. Values are mean ± s.d. of at least three experiments. **d**) Evolved lineages were analyzed by time-lapse microscopy for growth. The fraction of cells with extended lag-phase (white circles) or reduced growth rate (black circle) are shown for each day of the selection. **e)** The acquisition of a *nuoM* mutation in the *nuoN* hyper-tolerant strain background results in loss of tolerance. Survival to tobramycin (32 μg/ml) is scored over time for the strains indicated. Values are mean ± s.d. of at least n = 3 experiments. Data were normalized to CFUs at *t* = 0 hrs.

**Extended Data Figure 4:**
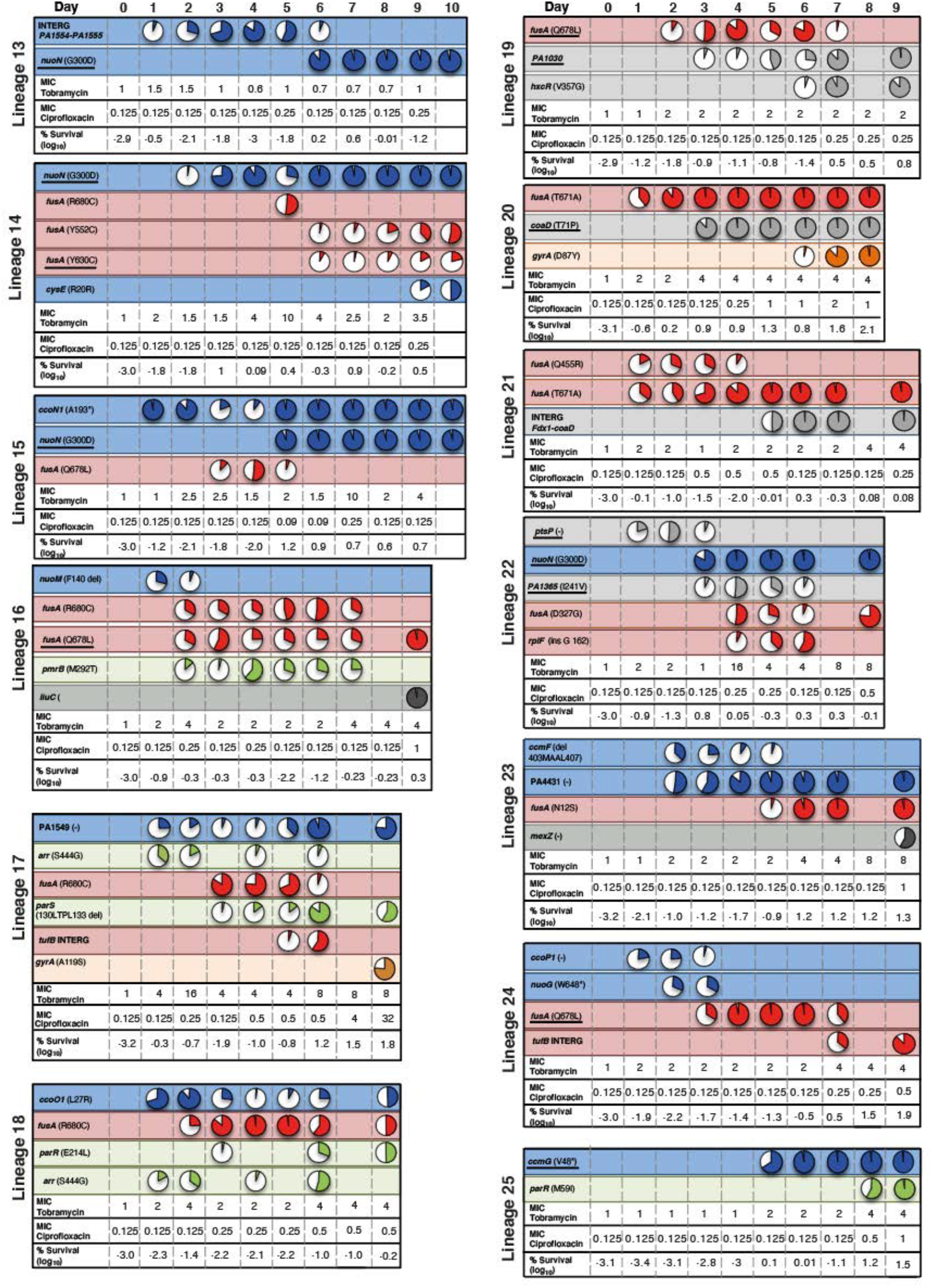
Mutations arising during combination drug selection as identified by whole genome sequencing. Lineage-specific mutations are indicated (see also Extended Table 1). Mutations in genes involved in respiration, protein synthesis, global regulators or other function are indicated in blue, red, green and grey, respectively. Circles indicate specific mutations with filled areas in the pie-charts indicating mutation distribution. For marked lineages (*) single colonies were randomly picked and allele combinations were analyzed by PCR. Specific mutations (underlined) were crossed back into the ancestral background and analyzed for their contribution to increased survival and MIC. MIC values and survival rates are reported for each day.

**Extended Data Figure 5:**
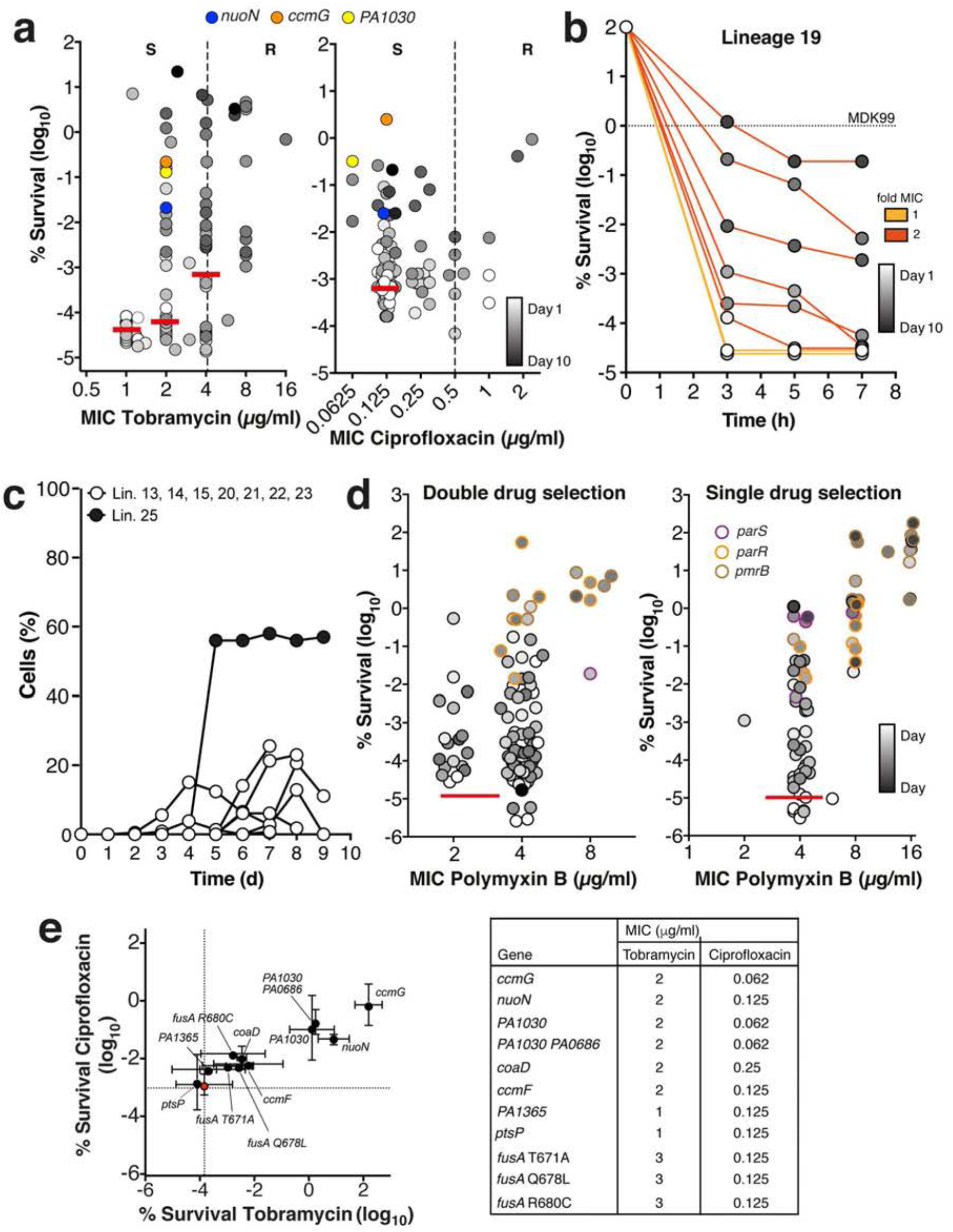
*P. aeruginosa* adaptation to double drug treatment. **a)** Tolerance levels and MIC of lineages evolved during selection with tobramycin (64 x MIC, left panel) or ciprofloxacin (80 x MIC, right panel). Strains generated by crossing back selected alleles (see Supplementary Table 1) into the ancestor are indicated in color. Average survival of the ancestor (n = 13) or lab strains with corresponding MICs below the clinical breakpoint of resistance ^30^ (dotted lines) is indicated by red lines. The dotted lines mark the breakpoints of resistance according to EUCAST: S, sensitive; R, resistant. **d)** Example of tolerance evolution in lineage 19. Samples from sequential time points (grey scale) were treated with tobramycin (64 μg/ml) and survival was measured over time. MIC of the evolved population are indicated by the colored lines. Dotted-line, minimal duration of killing for 99% of the population (MDK99). **c)** Time lapse analysis of lineages evolved in the presence of tobramycin and ciprofloxacin reveals cells with different growth behavior. The fraction of cells of individual lineages with a prolonged lag-phase (white circles) or with reduced growth rate (black circle) are shown as a function of time. **d)** Evolved lineages show cross tolerance against polymyxin B. Lineages evolved during single or double drug selection were analyzed for polymyxin B resistance (MIC) and for survival during 3 hrs of treatment with 15 μg/ml polymyxin B. The time of isolation (days) of individual strains during the selection window is indicated in grey shading. Colors indicate the presence of SNPs in genes known to confer polymyxin resistance. Red line, average survival of the ancestor (*n* = 14). Data were normalized to CFU at *t* = 0 hrs. **e)** Contribution of individual mutant alleles to antibiotic survival. Selected alleles from evolved strains (Extended data Figs. 2, 4) were crossed into the PAO1 ancestor and MIC values (right) and survival (left) of the resulting mutants were determined. Survival was determined after 3h of treatment with tobramycin 16 μg/ml (x-axis) or ciprofloxacin 2.5 μg/ml (y-axis). Survival scores of the ancestor are indicated by a red circle. Values are mean ± s.d. (n > 3).

**Extended Data Figure 6:**
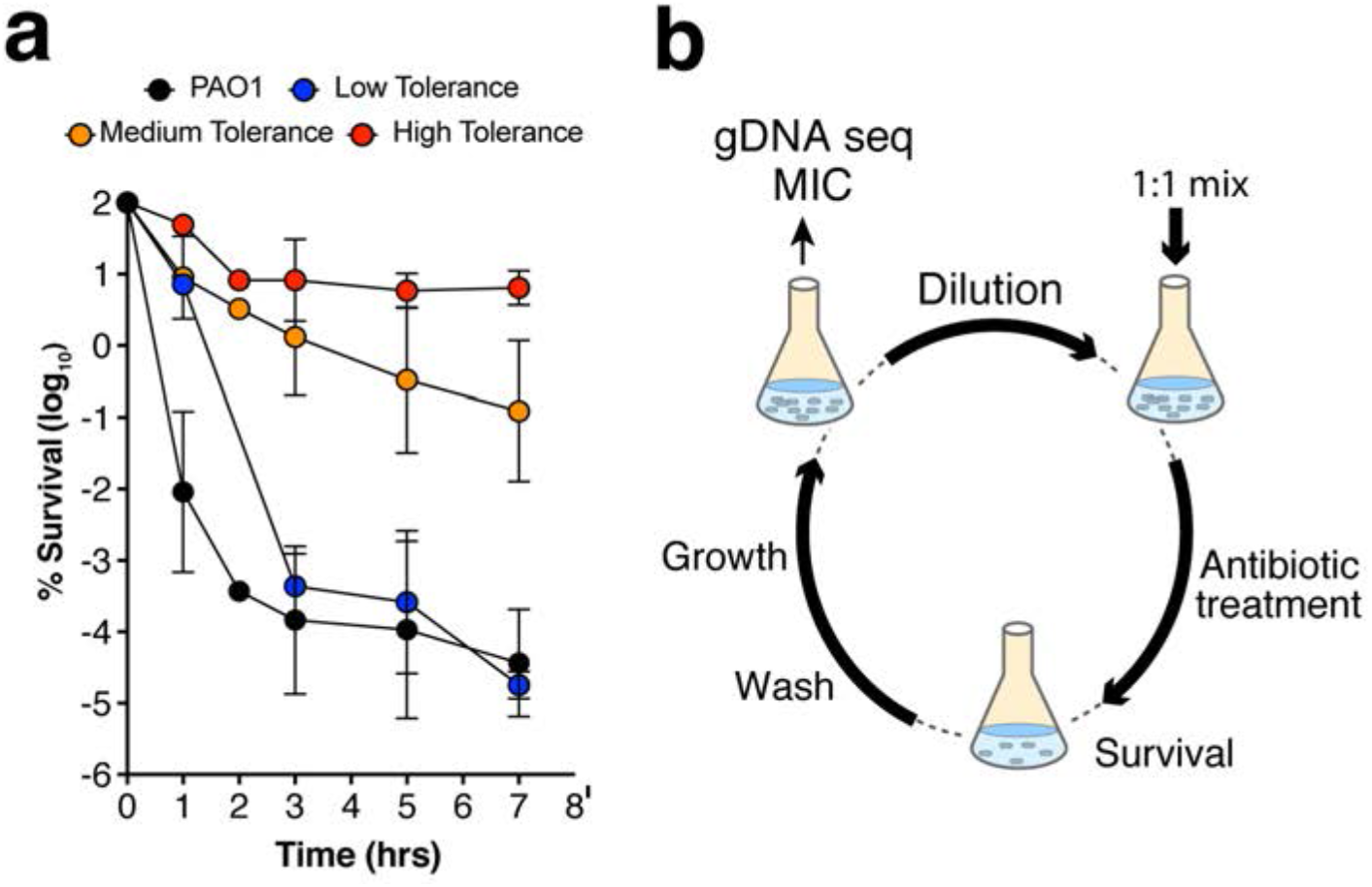
**a)** Mutant strains with different level of tolerance (wild type, low (nuoD_R551P_, medium, PA1030* and high, nuoN_G300D_) were treated with tobramycin (16 μg/ml) and survival was scored over time. Values are mean ± s.d. of at least three experiments. Data were normalized to CFU at *t* = 0 hours. **b**) Experimental design for iterative exposure of mixtures of *P. aeruginosa* strains with low and high tolerance levels to bactericidal antibiotics. Overnight cultures were diluted at a 1:1 ratio into fresh medium and challenged for 3 hrs with 32 μg/ml tobramycin. After antibiotic washout, cultures were re-suspended in fresh medium and grown over night. The cycle was repeated for several days until resistance increased. Whole population genome sequencing was performed for the ancestor mixture and the evolved lineages.

**Extended Data Figure 7:**
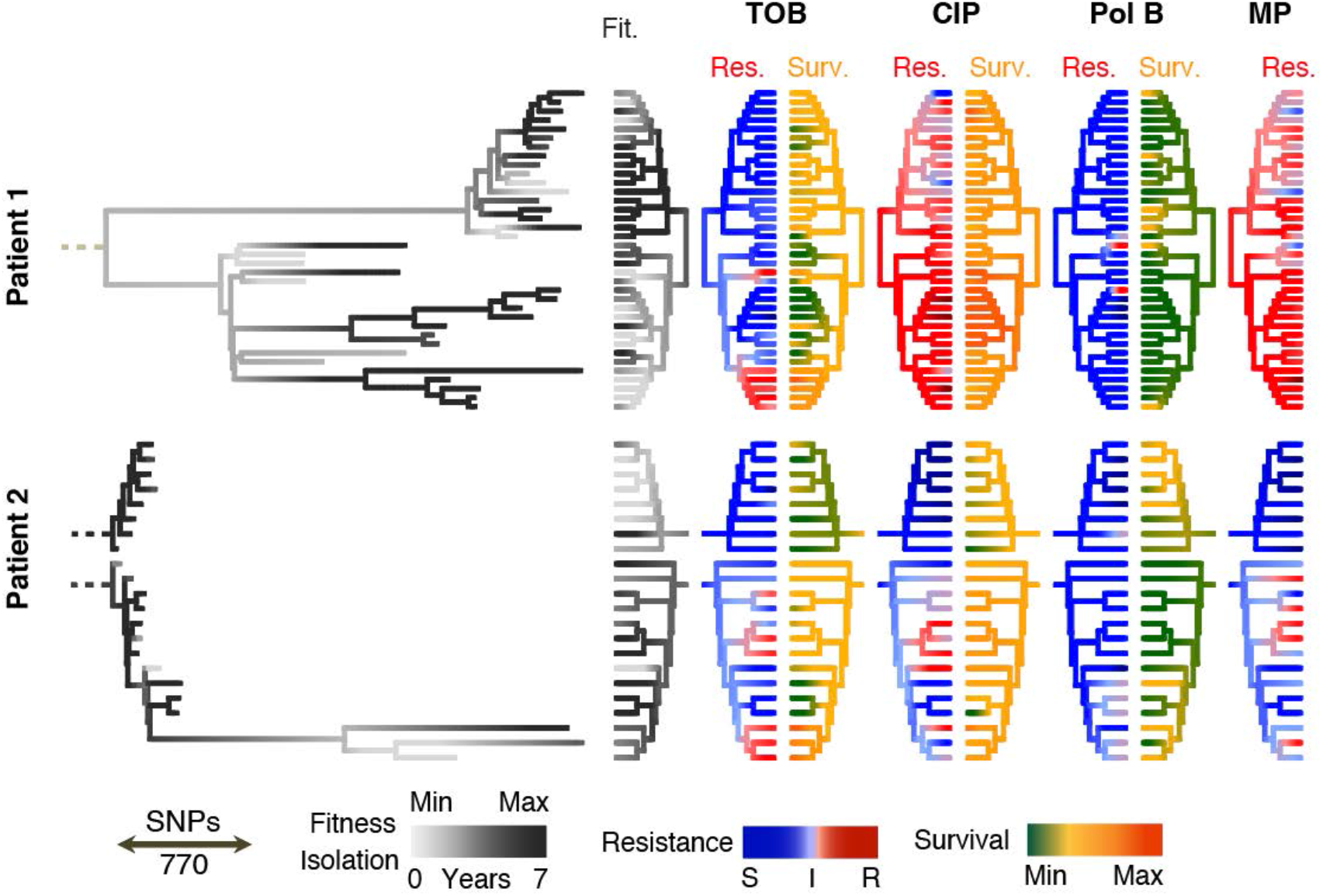
Micro-evolution of survival and antibiotic resistance during chronic infections of CF lungs. Genome-based phylogeny (85.579 positions) of *P. aeruginosa* strains from two CF patients sampled over a period of five years. Branch length of trees (left) indicate genetic differences (substitutions). Dashed roots represent extrapolated clonal ancestors. The colors of the terminal branches indicate (from left to right) isolation time during longitudinal sampling; fitness (growth in LB); resistance (Res) (average log2 fold variation from the intermediate resistance MIC of each antibiotic) and survival (Surv) (standardized survival scores for each antibiotic, see Material and Methods). Survival is reported for all strains including isolates with high MIC for a given antibiotic. TM, tobramycin; CIP, ciprofloxacin; PolB, polymyxin B, MP, meropenem.

**Extended Data Figure 8:**
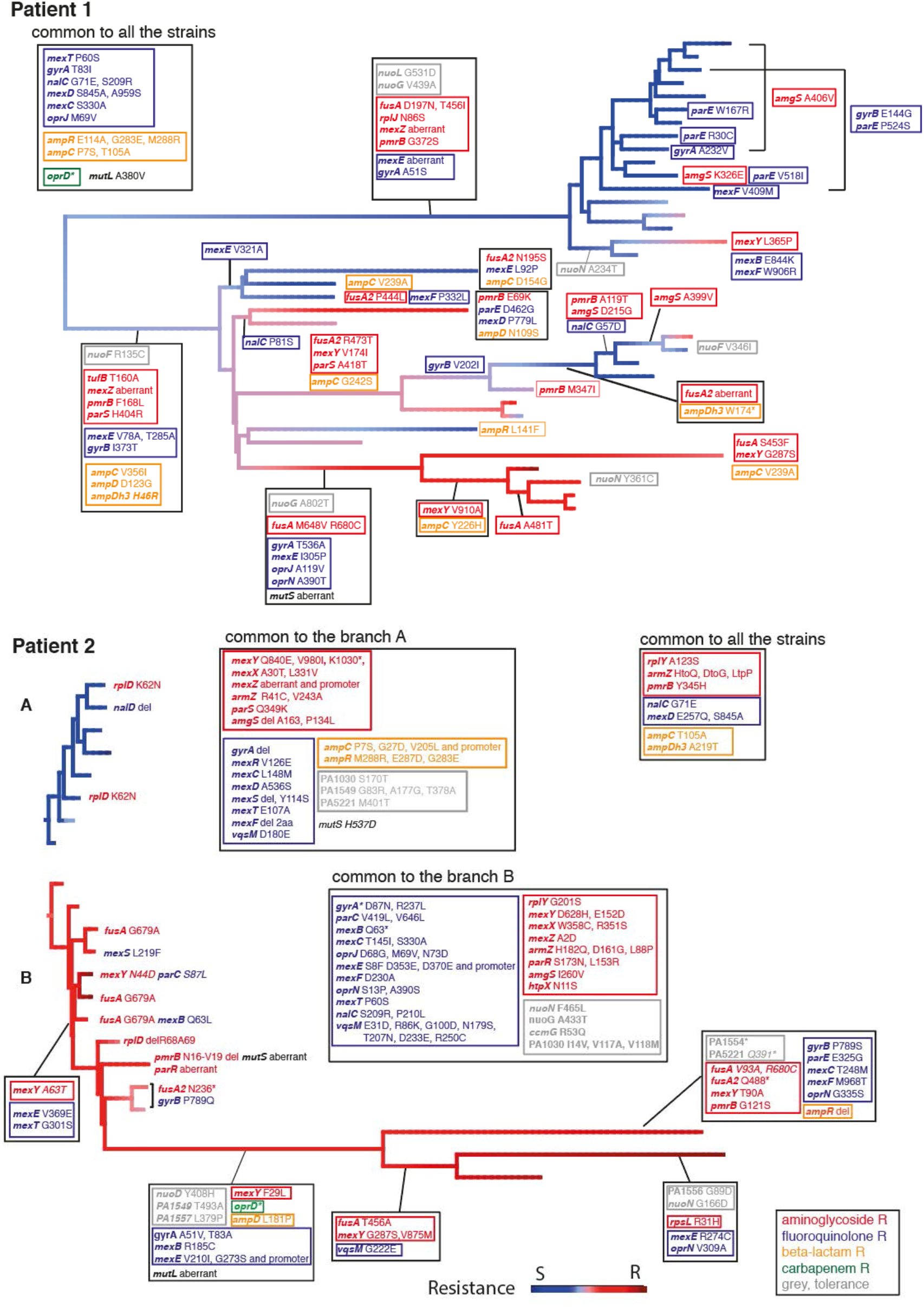
Alleles associated with antibiotic resistance and tolerance in *P. aeruginosa* isolates from CF patients. Genome based phylogeny of *P. aeruginosa* strains from two CF patients as shown in Fig. 5b. The colors of the terminal branches of the phylogeny indicate the resistance score as described in Fig. 5b. Mutations involved in resistance to aminoglycosides (red), fluoroquinolones (blue), carbapenems (green) and other β-lactams (orange) are indicated for each *P. aeruginosa* isolate from patient 1 and 2. Mutations in genes that were shown to contribute to tolerance in this study are highlighted in grey.

## Supplementary information

### Ethics statement

The clinical *P. aeruginosa* isolates used in this study were cultured from patient samples collected for routine microbiological testing at the University Hospital, Basel. Sub-culturing and analysis of bacteria were performed anonymously. No additional procedures were carried out on patients. Cultures were sampled following regular procedures with written informed consent, in agreement with the guidelines of the Ethikkommission beider Basel EKBB.

### Data availability statement

All genetic variations observed during *in vitro* evolution of *P. aeruginosa PAO1* are referenced in the Supplementary Table 1. Raw sequencing data of clinical strains will be deposited under an accession number available at publication time at the NCBI Sequence Read Archive (SRA).

## Supplementary Methods

### Bacterial strains and culture conditions

Strains used in this study are listed in Supplementary Table 2. Unless otherwise stated, *P. aeruginosa* PA01 and all *E. coli* strains were grown at 37°C in Luria Bertani (LB) medium ^1^ under shaking at 170 rpm or alternatively static and solidified with 1.3% agar when appropriate. For *P. aeruginosa*, tetracycline at 100 μg/ml (*E. coli* 12.5 μg/ml).

### Plasmids and oligonucleotides

Plasmid and primers used in this study are listed in Supplementary Table 2 and 3.

### Molecular biology procedures

Cloning was carried out in accordance with standard molecular biology techniques. Plasmids pEX18Tc-*nuoN*\*, pEX18Tc-*nuoD*\*, pEX18Tc-*nuoM*\*, pEX18Tc-*PA5221*\*, pEX18Tc-*ccmG*\* pEX18Tc-*coaD*\* pEX18Tc-*parS*\* pEX18Tc-*FusA*Y630C, pEX18Tc-*FusA*Q678L pEX18Tc-*PA1549*\* were produced by ligation of the *nuoN*\* (primer A and B, from Lineage 14, day 7 genomic DNA), *nuoD*\* (primer C and D, from Lineage 14, day 1 genomic DNA), *nuoM*\* (primer E and F, from Lineage 14, day 9 genomic DNA) *FusA* Y630C (primer G and H, from Lineage 2, day 10 genomic DNA), *FusA* Q678L (primer G and H, from Lineage 7, day 4 genomic DNA), *PA1549*\* (primer I and J, from Lineage 23, day 1 genomic DNA), *ccmG*\* (primer K and L, from Lineage 13, day 7 genomic DNA), *coaD*\* (primer M and N, from Lineage 13, day 5 genomic DNA), *parS*\* (amplified with primers O and P, from Lineage 18, day 5 genomic DNA) and *PA5221*\* (amplified with primers Q and R from Lineage 17, day 2 genomic DNA) PCR fragment between the *Hind*III and *Xba*I sites of pEX18Tc ^2^. The strains carrying the mutation *fusA* T671A, *fusA* R680C, *ccmF*\* and *PA1030*\* *PA0686*\* are single colony isolated from the lineage 8 (day 2), 6 (day 4), 24 (day 2) and 7 (day 7), respectively.

### Evolution experiments

Independent overnight cultures of PAO1 single colony in 5 ml of LB were diluted into fresh LB medium to OD 0.12 and challenged with antibiotics for 3h at 37°C, 170 rpm in flask. At indicated time points, aliquots of the cultures were sampled, diluted to the appropriate dilutions and plated on LB plates. CFUs were measured after overnight incubation and over a two-day period to make sure that all CFUs had appeared. After antibiotic treatment, cells were washed two times in LB to remove antibiotics and the pellet was inoculated into 5 ml of LB fresh medium for another by plating out before and after treatment. In the evolution with single drug, tobramycin was used at 32 μg/ml. In the evolution with two drugs, combination of tobramycin and ciprofloxacin at 16 μg/ml and 2.5 μg/ml, respectively was used. Every day, before the drug treatment: i) 1 ml of the overnight culture was pelleted and the pellet frozen at −80°C in 1 ml of LB 10% DMSO for further analysis; ii) 1 ml of the overnight culture was pelleted and the pellet frozen for genomic DNA extraction; iii) MIC of the population was estimated (see below for details). For competition experiments the followed protocol was the same except that the overnight cultures of the competing strains were diluted into 20 ml fresh LB medium to a final OD 0.06, each and challenge with 32 μg/ml tobramycin for 3h at 37°C, 170 rpm in flask. In order to assess the right ratio (1:1) of the competing strains, at day 0, before the drug treatment, 1 ml of the mixture was pelleted and the pellet frozen for genomic DNA extraction.

### Antibiotic survival assays

To measure the survival under antibiotic treatments, overnight cultures, each grown from a single colony in LB in case of the mutants or from 10% DMSO stock for the evolved lineages, were diluted to OD 0.12 into fresh LB medium supplemented with a fixed antibiotics concentration in order to avoid possible changes in the mode of action of the different drugs used. At indicated time points, aliquots of the cultures were sampled, diluted and plated on LB plates. CFUs were counted after overnight incubation and over a 2 days period to make sure that all CFUs had appeared.

### MIC assays

MICs were quantified based on adaptations of methods described elsewhere ^3,4^. Briefly, an overnight culture was diluted in LB to an inoculum of 1 × 10^6^ CFU / ml, incubated in a range of two-fold antibiotic dilutions and grown for 16–20 h at 37°C with shaking. After incubation, the OD at 600 nm (OD_600_) was measured. We defined the MIC value as the lowest antibiotic concentration where no growth was observed

### Microscopy

Bacteria were grown over night (ON) (OD_600_ 2-3) in LB liquid medium, diluted 1:100 in fresh LB and then transferred to a slide with a 1% agarose pad buffered with LB medium. Bacteria were imaged by time-lapse microscopy using a DeltaVision microscope (Applied Precision) equipped with a 100× oil immersion objective and an environmental chamber maintained at 35°C. Images were recorded on phase-contrast using a CoolSnap HQ2 camera. Images were processed using Softworx software (Applied Precision). Lag-phase was defined as the time between the start of the movie and the initiation of the cell elongation.

### Genomic DNA extraction, whole-genome sequencing and coverage analyses

For each evolved lineage 1 ml of the ON culture was daily pelleted and frozen at −80°C. Clinical isolates of *P. aeruginosa* were grown ON in liquid LB and pellets from 1 ml of culture were collected. Genomic DNA was then extracted using the GenElute Bacterial Genomic DNA kit (Sigma, NA2120-1KT). Genomic DNA integrity was monitored by 0.6% agar gel electrophoresis. And gDNA was send to the Genomics Facility Basel for library preparation and Illumina sequencing (ETH Zurich Department of Biosystems Science and Engineering, Basel, Switzerland). Library preparation and sample barcoding were generated with the Nextera XT approach (Illumina) and libraries quality was checked with a Fragment Analyzer (Advanced Analytical). PE125 sequencing runs were performed on 95 libraries at the time on a single HiSeq lane (Illumina) with a targeted coverage of *c.a.* 100X. Reads were mapped onto the genome of the reference strain *Pseudomonas aeruginosa PAO1* (NC_002516) with Bowtie 2 ^5^ and small polymorphisms, structural and coverage variants were spotted with Samtools ^6^ and a collection of in-house Perl scripts. Only genetic variations that were observed in more than 10% of the reads in at least one time point are reported here. In many cases, the evolutionary context was simple enough to infer the presence of sub-lineages partially or fully genetically identifiable by coverage analysis of the population genome sequencing. For example, in lineage 1 (Extended fig 2), one can see the sub-population exhibiting an intergenic mutation between PA1554 and PA1555 invading the lineage over the five first days while the number of wild type variants drops. At day six, the wild type variant of the intergenic region become dominant again in the evolving population. At the same time and with similar levels, another variation in the nuoN gene quickly invaded the lineage indicating that these two mutations represent two competing sub-lineages. Furthermore, often, genetic fixations could be observed in many lineages when 100% of sequencing reads exhibited a stable mutation over several time points (e.g. lineages 2, 8-11, 13-19, 21-23, 25). In these cases, we could establish sequential acquisition of genetic variations (when present in more than 10% of the population).

### Phylogeny of clinical strains

Whole genome sequences of 58 chronic infection isolates from patients one and two were compared to references from GenBank revealing highest similarities with strain RIVM-EMC2982, a clinical isolate from the Netherlands (CP016955.1), 12-4-4(59), a strain isolated from blood of a burn patient (NZ_CP013696.1) and W36662, a strain isolated from a cancer patient (CP008870.2). Similarity to the references was assessed by the similarity Blast Score (in bits) of largest contigs from each assembly. The phylogenetic tree of the chronic infection strains from patients one and two were based on the 85.579 chromosomal positions where polymorphism has been observed in at least one of the clinical strains sequenced. The evolutionary history was inferred in MEGA7 ^7^ by using the Maximum Likelihood method based on the Tamura-Nei model ^8^. Phenotypically color-coded trees were generated with the R-package phytools (http://github.com/liamrevell/phytools). Differences between the clonal branches (i.e. clades closely related to the same reference) were between 17,400 and 27,400 base pairs at conserved sites (including polymorphisms and indels).

### Computation of Tolerance and Resistance scores, clustering and surface fitting over time

For longitudinal samples isolated from single patients (Fig. 4a), resistance score (R) were computed as the average of log2 fold variations from the intermediary clinical resistance level (I) for each drug considered (d): tobramycin (I: 6 μg/ml), ciprofloxacin (I: 1 μg/ml), polymyxin B (I: 4 μg/ml), colistin (I: 4 μg/ml), meropenem (I: 6 μg/ml). Therefore, for a given strain considering five drugs,

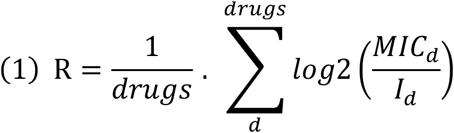

Tolerance scores (*T*) correspond to the average of log10 values for a three hours survival assay (SURV, see here above) with different drugs (*d*) and standardized per patient (*pat*). Only survival values from non-resistant strains were considered to compute the latter tolerance score with MIC cut-off that minimized the impact of resistance traits on the survival assay: tobramycin (I: 2 μg/ml), ciprofloxacin (I: 1 μg/ml), polymyxin B (I: 8 μg/ml). Therefore, for a given strain considering one to three drugs,

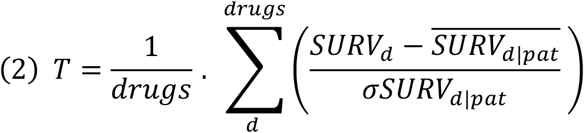

For the analysis of tolerance and resistance over 317 isolates (Fig. 4b), the resistance scores correspond to the MIC for tobramycin or ciprofloxacin normalized to the clinically relevant resistance break points similarly to equation (1). Tolerance scores correspond to log10 values for a 3 hours survival assay (see above) with either tobramycin or ciprofloxacin standardized by drug over the whole data set similarly to equation (2) but with average and standard deviation from the whole strain data set. In order to assess resistance and tolerance of the same strains, we only report tolerance score for strains that were sensitive to at least one of the two drugs, tobramycin (only strains with MIC <4 μg/ml) or ciprofloxacin (only strains with MIC ≤1 μg/ml) and we provide tolerance score for drug different to the one used to assess resistance. This made it possible to gauge the multidrug tolerance feature under at least one effective antimicrobial treatment. Visible structure in the data suggested the existence of different resistance/tolerance profiles over microevolution (Extended Data Fig. 9). Indeed, iterative clustering around medoids (R, package: cluster, function: pam) identified the optimal (i.e. smallest) number of clusters (i.e. 3) that best minimized the intra-group heterogeneity (Elbow method). Clusters are shown on the principal component analysis plot of tolerance, resistance and age of patient at isolation time. Surface fitting of tolerance and resistance scores of strains over the age of patients at isolation time was performed with the Tps function of the Fields R package. This function performs a thin plate spline regression as method of interpolation with smoothing parameters automatically chosen by generalized cross-validation. Naïve isolated were used to simulate the primo-infecting strains and were assigned an age of patient of zero. To avoid useless interpolations, regions with no or poor data coverage were excluded. Clustering of killing profiles was performed with the “cluster” R library using the pam (Partitioning Around Medoids) function. Appropriate numbers of clusters were selected by the “elbow method” on Ball & Hall sores computed with the intCriteria function from the clusterCrit R library.

### Statistics

Data acquired were analyzed using GraphPad Prism version 6.03 for Mac and R.3.5.0 with different packages as described in appropriate sections. Mean values and standard deviations were obtained from at least three independent experiments (biological replicates). All the documented results are highly reproducible. No statistical method was used to predetermine the sample size.

---

**Supplementary Table 2.**
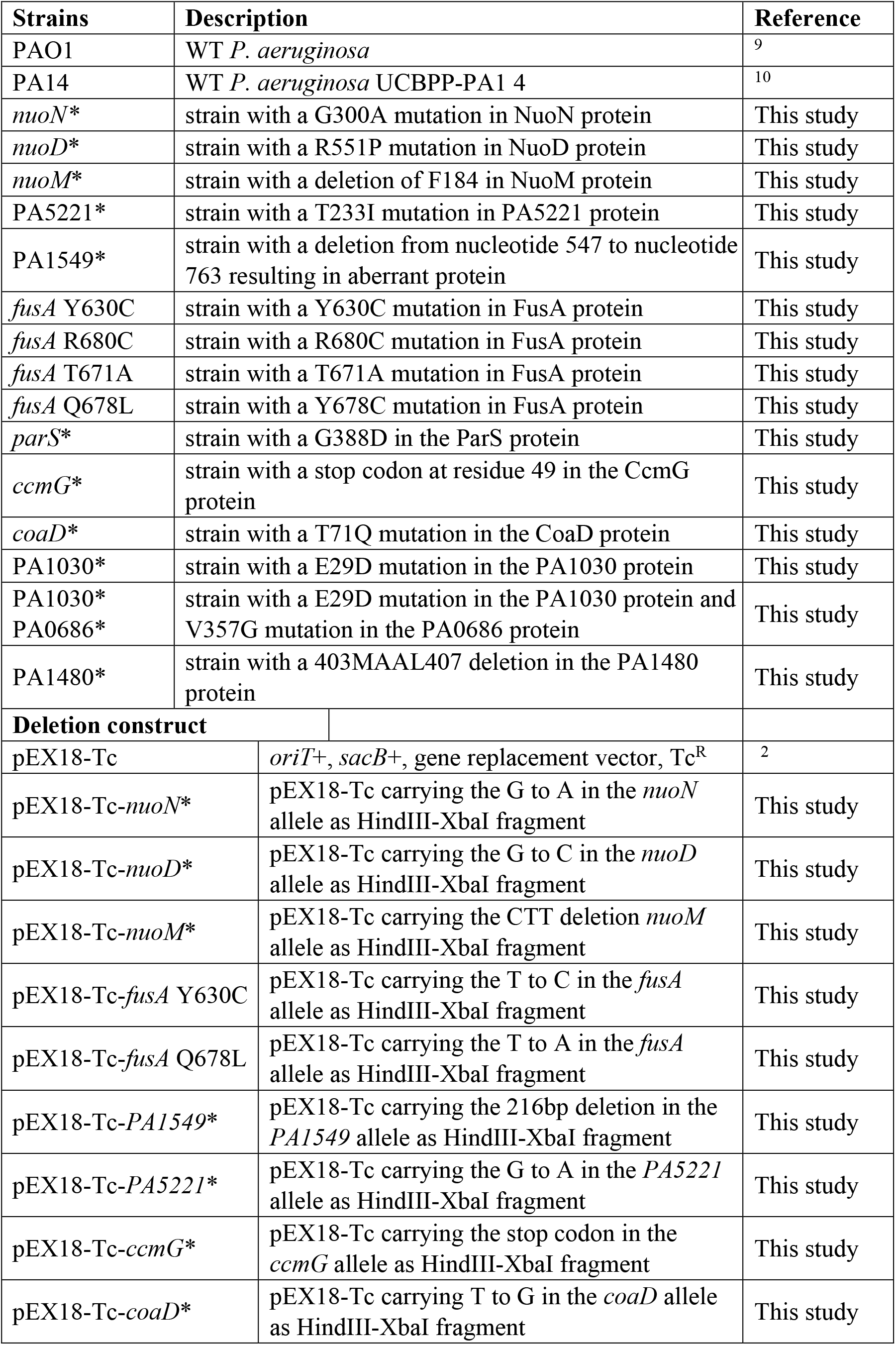
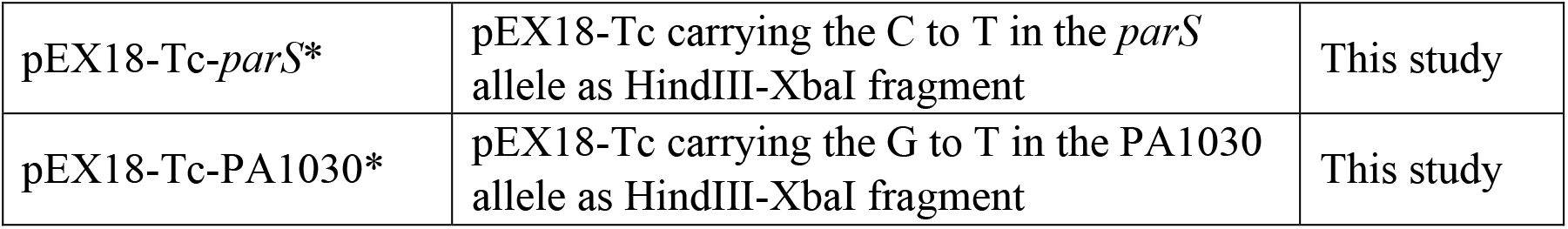
List of strains and plasmids used in this study.

**Supplementary Table 3.**
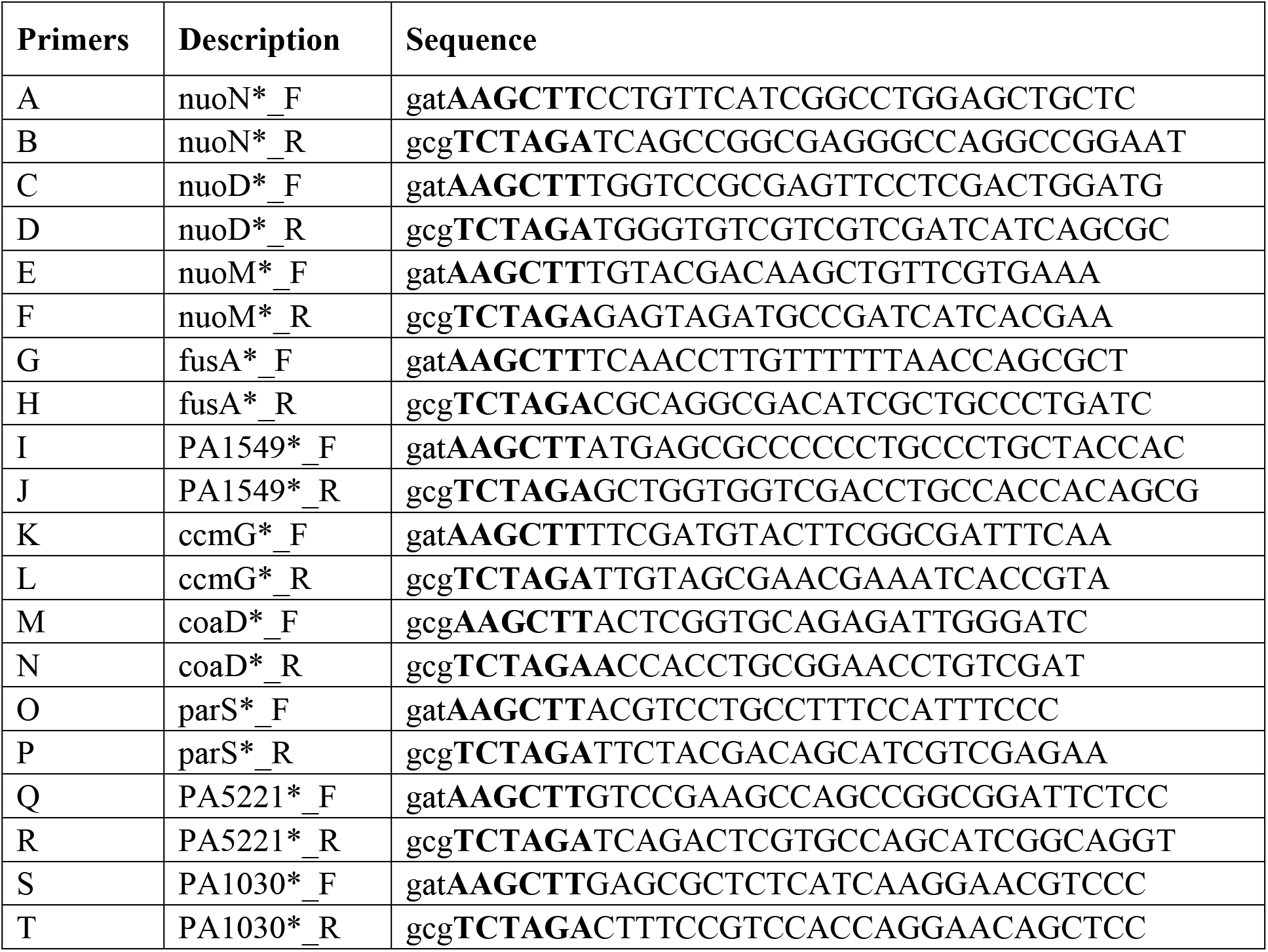
Oligos used in this study.

**Table S1.**
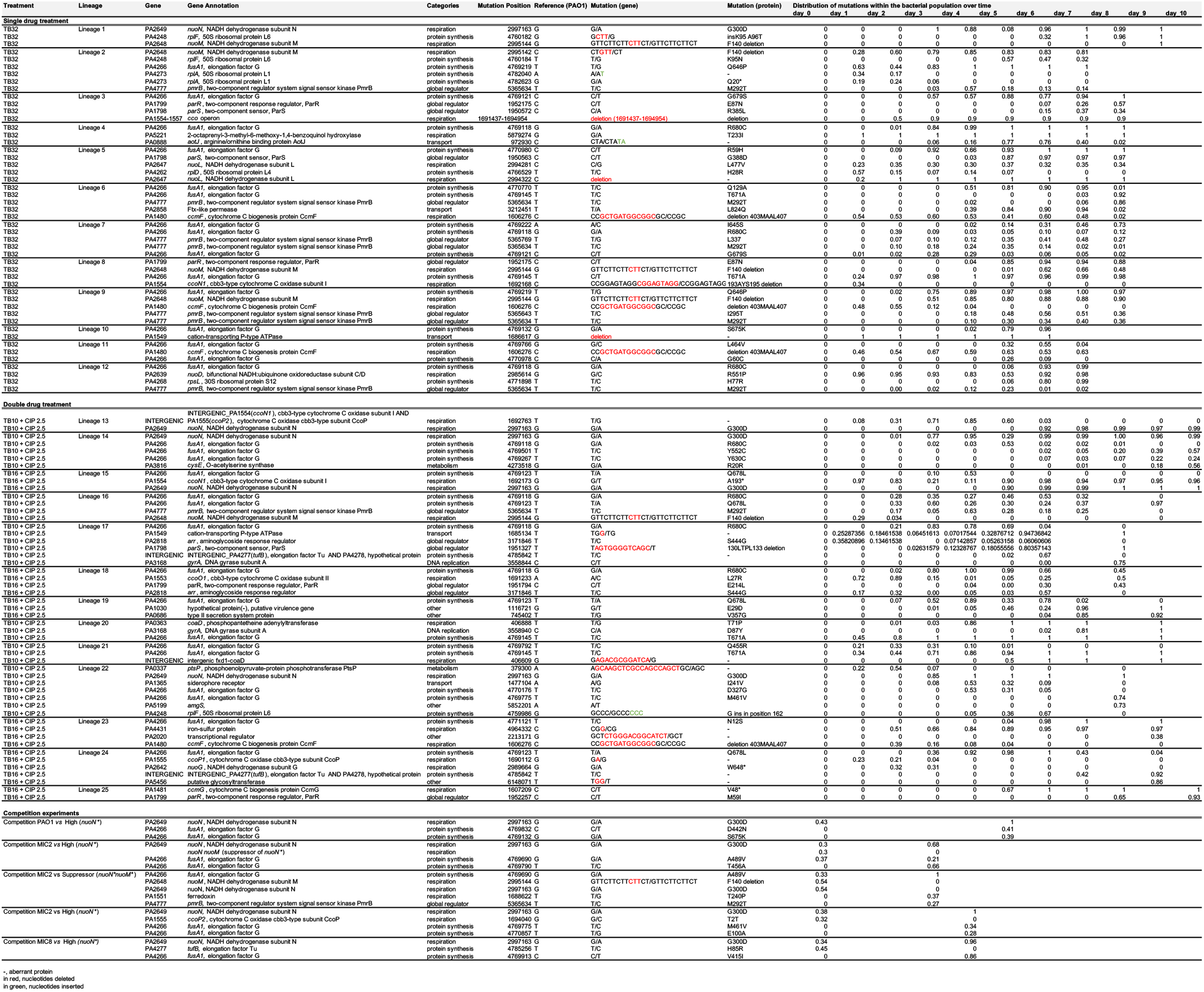
List of mutations acquired during the evolution with single or double drug treatment

## References

1 Hajjeh, R. a. M., A. Global action plan on antibiotic resistance. (WHO, 2016).

2 Andersson, D. I. et al. Antibiotic resistance: turning evolutionary principles into clinical reality. FEMS Microbiol Rev 44, 171–188, doi:10.1093/femsre/fuaa001 (2020).

3 Horne, D. & Tomasz, A. Tolerant response of *Streptococcus sanguis* to beta-lactams and other cell wall inhibitors. Antimicrob Agents Chemother 11, 888–896 (1977).

4 Lewis, K. The Science of Antibiotic Discovery. Cell 181, 29–45, doi:10.1016/j.cell.2020.02.056 (2020).

5 Scholar, E. M. & Pratt, W. B. The antimicrobial drugs. 2nd edn, (Oxford University Press, 2000).

6 Brauner, A., Shoresh, N., Fridman, O. & Balaban, N. Q. An Experimental Framework for Quantifying Bacterial Tolerance. Biophys J 112, 2664–2671, doi:10.1016/j.bpj.2017.05.014 (2017).

7 Brauner, A., Fridman, O., Gefen, O. & Balaban, N. Q. Distinguishing between resistance, tolerance and persistence to antibiotic treatment. Nat Rev Microbiol 14, 320–330, doi:10.1038/nrmicro.2016.34 (2016).

8 Bigger, J. W. Treatment of staphylococcal infections with penicillin by intermittent sterilisation. Lancet 244, 497–500 (1944).

9 Fridman, O., Goldberg, A., Ronin, I., Shoresh, N. & Balaban, N. Q. Optimization of lag time underlies antibiotic tolerance in evolved bacterial populations. Nature 513, 418–421, doi:10.1038/nature13469 (2014).

10 Van den Bergh, B. et al. Frequency of antibiotic application drives rapid evolutionary adaptation of *Escherichia coli persistence*. Nat Microbiol 1, 16020, doi:10.1038/nmicrobiol.2016.20 (2016).

11 Liu, J., Gefen, O., Ronin, I., Bar-Meir, M. & Balaban, N. Q. Effect of tolerance on the evolution of antibiotic resistance under drug combinations. Science 367, 200–204, doi:10.1126/science.aay3041 (2020).

12 Windels, E. M., Van den Bergh, B. & Michiels, J. Bacteria under antibiotic attack: Different strategies for evolutionary adaptation. PLoS Pathog 16, e1008431, doi:10.1371/journal.ppat.1008431 (2020).

13 Jorth, P. et al. Regional Isolation Drives Bacterial Diversification within Cystic Fibrosis Lungs. Cell Host Microbe 18, 307–319, doi:10.1016/j.chom.2015.07.006 (2015).

14 Ramsey, B. W. et al. Intermittent administration of inhaled tobramycin in patients with cystic fibrosis. Cystic Fibrosis Inhaled Tobramycin Study Group. N Engl J Med 340, 23–30, doi:10.1056/NEJM199901073400104 (1999).

15 Tuomanen, E., Durack, D. T. & Tomasz, A. Antibiotic tolerance among clinical isolates of bacteria. Antimicrob Agents Chemother 30, 521–527 (1986).

16 Hofsteenge, N., van Nimwegen, E. & Silander, O. K. Quantitative analysis of persister fractions suggests different mechanisms of formation among environmental isolates of *E. coli*. BMC Microbiol 13, 25, doi:10.1186/1471-2180-13-25 (2013).

17 Mulcahy, L. R., Burns, J. L., Lory, S. & Lewis, K. Emergence of *Pseudomonas aeruginosa* strains producing high levels of persister cells in patients with cystic fibrosis. J Bacteriol 192, 6191–6199, doi:10.1128/JB.01651-09 (2010).

18 Stepanyan, K. et al. Fitness trade-offs explain low levels of persister cells in the opportunistic pathogen *Pseudomonas aeruginosa*. Mol Ecol 24, 1572–1583, doi:10.1111/mec.13127 (2015).

19 Meylan, S., Andrews, I. W. & Collins, J. J. Targeting Antibiotic Tolerance, Pathogen by Pathogen. Cell 172, 1228–1238, doi:10.1016/j.cell.2018.01.037 (2018).

20 Levin-Reisman, I. et al. Antibiotic tolerance facilitates the evolution of resistance. Science 355, 826–830, doi:10.1126/science.aaj2191 (2017).

21 Kester, J. C. & Fortune, S. M. Persisters and beyond: mechanisms of phenotypic drug resistance and drug tolerance in bacteria. Crit Rev Biochem Mol Biol 49, 91–101, doi:10.3109/10409238.2013.869543 (2014).

22 Scotet, V., L’Hostis, C. & Ferec, C. The Changing Epidemiology of Cystic Fibrosis: Incidence, Survival and Impact of the CFTR Gene Discovery. Genes (Basel) 11, 589, doi:10.3390/genes11060589 (2020).

23 Hansen, C. R., Pressler, T. & Hoiby, N. Early aggressive eradication therapy for intermittent *Pseudomonas aeruginosa* airway colonization in cystic fibrosis patients: 15 years experience. J Cyst Fibros 7, 523–530, doi:10.1016/j.jcf.2008.06.009 (2008).

24 La Rosa, R., Johansen, H. K. & Molin, S. Convergent Metabolic Specialization through Distinct Evolutionary Paths in *Pseudomonas aeruginosa*. MBio 9, e00269–00218, doi:10.1128/mBio.00269-18 (2018).

25 Speert, D. P. et al. Epidemiology of *Pseudomonas aeruginosa* in cystic fibrosis in British Columbia, Canada. Am J Respir Crit Care Med 166, 988–993 (2002).

26 Fothergill, J. L., Walshaw, M. J. & Winstanley, C. Transmissible strains of *Pseudomonas aeruginosa* in cystic fibrosis lung infections. Eur Respir J 40, 227–238, doi:10.1183/09031936.00204411 (2012).

27 Jelsbak, L. et al. Molecular epidemiology and dynamics of *Pseudomonas aeruginosa* populations in lungs of cystic fibrosis patients. Infect Immun 75, 2214–2224, doi:10.1128/IAI.01282-06 (2007).

28 Cramer, N., Wiehlmann, L. & Tummler, B. Clonal epidemiology of *Pseudomonas aeruginosa* in cystic fibrosis. Int J Med Microbiol 300, 526–533, doi:10.1016/j.ijmm.2010.08.004 (2010).

29 Langton Hewer, S. C. & Smyth, A. R. Antibiotic strategies for eradicating *Pseudomonas aeruginosa* in people with cystic fibrosis. Cochrane Database Syst Rev 4, CD004197, doi:10.1002/14651858.CD004197.pub5 (2017).

30 EUCAST. Breakpoint tables for interpretation of MICs and zone diameters. European Committee on Antimicrobial Susceptibility Testing Version 8.1, 2018.(2018).

31 Geller, D. E., Pitlick, W. H., Nardella, P. A., Tracewell, W. G. & Ramsey, B. W. Pharmacokinetics and bioavailability of aerosolized tobramycin in cystic fibrosis. Chest 122, 219–226 (2002).

32 Bolard, A., Plesiat, P. & Jeannot, K. Mutations in Gene *fusA1* as a Novel Mechanism of Aminoglycoside Resistance in Clinical Strains of *Pseudomonas aeruginosa*. Antimicrob Agents Chemother 62, e01835–01817, doi:10.1128/AAC.01835-17 (2018).

33 Tepekule, B., Uecker, H., Derungs, I., Frenoy, A. & Bonhoeffer, S. Modeling antibiotic treatment in hospitals: A systematic approach shows benefits of combination therapy over cycling, mixing, and mono-drug therapies. PLoS Comput Biol 13, e1005745, doi:10.1371/journal.pcbi.1005745 (2017).

34 Verstraeten, N. et al. Obg and Membrane Depolarization Are Part of a Microbial Bet-Hedging Strategy that Leads to Antibiotic Tolerance. Mol Cell 59, 9–21, doi:10.1016/j.molcel.2015.05.011 (2015).

35 Freire, D. M. et al. An NAD(+) Phosphorylase Toxin Triggers *Mycobacterium tuberculosis* Cell Death. Mol Cell 73, 1282–1291 e1288, doi:10.1016/j.molcel.2019.01.028 (2019).

36 Skjerning, R. B., Senissar, M., Winther, K. S., Gerdes, K. & Brodersen, D. E. The RES domain toxins of RES-Xre toxin-antitoxin modules induce cell stasis by degrading NAD+. Mol Microbiol 111, 221–236, doi:10.1111/mmi.14150 (2019).

37 Piscotta, F. J., Jeffrey, P. D. & Link, A. J. ParST is a widespread toxin-antitoxin module that targets nucleotide metabolism. Proc Natl Acad Sci U S A 116, 826–834, doi:10.1073/pnas.1814633116 (2019).

38 Higgins, P. G., Fluit, A. C., Milatovic, D., Verhoef, J. & Schmitz, F. J. Mutations in GyrA, ParC, MexR and NfxB in clinical isolates of *Pseudomonas aeruginosa*. Int J Antimicrob Agents 21, 409–413 (2003).

39 Ahmad, M. H., Rechenmacher, A. & Bock, A. Interaction between aminoglycoside uptake and ribosomal resistance mutations. Antimicrob Agents Chemother 18, 798–806 (1980).

40 Fernandez, L. et al. Adaptive resistance to the “last hope” antibiotics polymyxin B and colistin in *Pseudomonas aeruginosa* is mediated by the novel two-component regulatory system ParR-ParS. Antimicrob Agents Chemother 54, 3372–3382, doi:10.1128/AAC.00242-10 (2010).

41 Loveland, A. B. et al. Ribosome*RelA structures reveal the mechanism of stringent response activation. eLlife 5, e17029, doi:10.7554/eLife.17029 (2016).

42 Macvanin, M., Johanson, U., Ehrenberg, M. & Hughes, D. Fusidic acid-resistant EF-G perturbs the accumulation of ppGpp. Mol Microbiol 37, 98–107 (2000).

43 Efremov, R. G., Baradaran, R. & Sazanov, L. A. The architecture of respiratory complex I. Nature 465, 441–445, doi:10.1038/nature09066 (2010).

44 Taber, H. W., Mueller, J. P., Miller, P. F. & Arrow, A. S. Bacterial uptake of aminoglycoside antibiotics. Microbiol Rev 51, 439–457 (1987).

45 Blokhina, S. V., Sharapova, A. V., Ol’khovich, M. V., Volkova capital Te, C. & Perlovich, G. L. Solubility, lipophilicity and membrane permeability of some fluoroquinolone antimicrobials. Eur J Pharm Sci 93, 29–37, doi:10.1016/j.ejps.2016.07.016 (2016).

46 Claudi, B. et al. Phenotypic variation of *Salmonella* in host tissues delays eradication by antimicrobial chemotherapy. Cell 158, 722–733, doi:10.1016/j.cell.2014.06.045 (2014).

47 Marvig, R. L., Sommer, L. M., Molin, S. & Johansen, H. K. Convergent evolution and adaptation of *Pseudomonas aeruginosa* within patients with cystic fibrosis. Nat Genet 47, 57–64, doi:10.1038/ng.3148 (2015).

48 Karna, S. L. et al. Genome Sequence of a Virulent *Pseudomonas aeruginosa* Strain, 12-4-4(59), Isolated from the Blood Culture of a Burn Patient. Genome Announc 4, e00079–00016, doi:10.1128/genomeA.00079-16 (2016).

49 Greipel, L. et al. Molecular Epidemiology of Mutations in Antimicrobial Resistance Loci of *Pseudomonas aeruginosa* Isolates from Airways of Cystic Fibrosis Patients. Antimicrob Agents Chemother 60, 6726–6734, doi:10.1128/AAC.00724-16 (2016).

50 Bartell, J. A. et al. Evolutionary highways to persistent bacterial infection. Nat Commun 10, 629, doi:10.1038/s41467-019-08504-7 (2019).

51 Stefani, S. et al. Relevance of multidrug-resistant *Pseudomonas aeruginosa* infections in cystic fibrosis. Int J Med Microbiol 307, 353–362, doi:10.1016/j.ijmm.2017.07.004 (2017).

52 Windels, E. M., Michiels, J. E., Van den Bergh, B., Fauvart, M. & Michiels, J. Antibiotics: Combatting Tolerance To Stop Resistance. mBio 10, e02095–02019, doi:10.1128/mBio.02095-19 (2019).

53 Kirby, A. E., Garner, K. & Levin, B. R. The relative contributions of physical structure and cell density to the antibiotic susceptibility of bacteria in biofilms. Antimicrob Agents Chemother 56, 2967–2975, doi:10.1128/AAC.06480-11 (2012).

54 Schurek, K. N. et al. Novel genetic determinants of low-level aminoglycoside resistance in *Pseudomonas aeruginosa*. Antimicrob Agents Chemother 52, 4213–4219, doi:10.1128/AAC.00507-08 (2008).

55 Frimodt-Moller, J. et al. Mutations causing low level antibiotic resistance ensure bacterial survival in antibiotic-treated hosts. Sci Rep 8, 12512, doi:10.1038/s41598-018-30972-y (2018).

56 Goldstein, F. The potential clinical impact of low-level antibiotic resistance in *Staphylococcus aureus*. J Antimicrob Chemother 59, 1–4, doi:10.1093/jac/dkl429 (2007).

57 Wistrand-Yuen, E. et al. Evolution of high-level resistance during low-level antibiotic exposure. Nat Commun 9, 1599, doi:10.1038/s41467-018-04059-1 (2018).

58 Conlon, B. P. et al. Activated ClpP kills persisters and eradicates a chronic biofilm infection. Nature 503, 365–370, doi:10.1038/nature12790 (2013).

59 Berney, M., Weilenmann, H. U. & Egli, T. Flow-cytometric study of vital cellular functions in *Escherichia coli* during solar disinfection (SODIS). Microbiology (Reading) 152, 1719–1729, doi:10.1099/mic.0.28617-0 (2006).

## Supplementary References

1 Miller, J. H. Experiments in molecular genetics. (Cold Spring Harbor Laboratory, 1972).

2 Hoang, T. T., Karkhoff-Schweizer, R. R., Kutchma, A. J. & Schweizer, H. P. A broad-host-range Flp-FRT recombination system for site-specific excision of chromosomally-located DNA sequences: application for isolation of unmarked *Pseudomonas aeruginosa* mutants. Gene 212, 77–86 (1998).

3 Wiegand, I., Hilpert, K. & Hancock, R. E. Agar and broth dilution methods to determine the minimal inhibitory concentration (MIC) of antimicrobial substances. Nat Protoc 3, 163–175, doi:10.1038/nprot.2007.521 (2008).

4 Liebens, V. et al. Identification and characterization of an anti-pseudomonal dichlorocarbazol derivative displaying anti-biofilm activity. Bioorg Med Chem Lett 24, 5404–5408, doi:10.1016/j.bmcl.2014.10.039 (2014).

5 Langmead, B. & Salzberg, S. L. Fast gapped-read alignment with Bowtie 2. Nat Methods 9, 357–359, doi:10.1038/nmeth.1923 (2012).

6 Li, H. A statistical framework for SNP calling, mutation discovery, association mapping and population genetical parameter estimation from sequencing data. Bioinformatics 27, 2987–2993, doi:10.1093/bioinformatics/btr509 (2011).

7 Kumar, S., Stecher, G. & Tamura, K. MEGA7: Molecular Evolutionary Genetics Analysis Version 7.0 for Bigger Datasets. Mol Biol Evol 33, 1870–1874, doi:10.1093/molbev/msw054 (2016).

8 Tamura, K. & Nei, M. Estimation of the number of nucleotide substitutions in the control region of mitochondrial DNA in humans and chimpanzees. Mol Biol Evol 10, 512–526, doi:10.1093/oxfordjournals.molbev.a040023 (1993).

9 Holloway, B. W. Genetic recombination in *Pseudomonas aeruginosa*. J Gen Microbiol 13, 572–581, doi:10.1099/00221287-13-3-572 (1955).

10 Rahme, L. G. et al. Common virulence factors for bacterial pathogenicity in plants and animals. Science 268, 1899–1902 (1995).

